# Episodic memory rescues working memory via pattern separation, pattern completion, and predictive recall

**DOI:** 10.64898/2026.08.26.746101

**Authors:** Zihan Bai, Daryl Fougnie, Sebastian Michelmann

**Affiliations:** Department of Psychology, New York University, New York.; Department of Psychology, New York University Abu Dhabi, Abu Dhabi.

## Abstract

Working memory is capacity-limited, but interactions with episodic memory may offset this constraint. We tested moment-by-moment contributions of episodic representations to working memory by combining the N-back and Mnemonic Similarity tasks. Thirty-one participants, undergoing eye-tracking, first encoded items in a one-back task, classifying them as “same” or “similar” to their predecessor. In a subsequent two-back task, mnemonic discrimination showed a graded, item-specific benefit of prior experience: performance was best for previously compared items, whereas recognition of identical repeats was unaffected. Successful discrimination of previously compared items was accompanied by greater pupil dilation, gradually emerging gaze patterns resembling those elicited by their similar pair-mate, and higher gaze-similarity between one-back and two-back target viewing. Diverging gaze patterns between pair-mates during one-back further predicted two-back discrimination. These findings challenge working memory’s characterization as an isolated system, demonstrating how it recruits episodic computations — encoding distinct traces, predicting upcoming content, and reinstating it at retrieval.

## Main

Humans routinely discriminate among similar objects, for instance, when picking out the right medication among nearly identical bottles – a task that requires maintaining, comparing, and updating recent information. Working memory (WM) has conventionally been credited as solely supporting this process, holding task-relevant information over short intervals via active neural maintenance of representations [1–4]. However, WM is capacity-limited and its representations degrade rapidly [5], a real constraint when discrimination depends on retaining fine-grained detail about several similar items at once. In everyday experience, where information arrives in rapid succession, more durable memory traces may be needed to support discrimination when WM capacity is exceeded.

Episodic memory (EM) is a natural candidate to afford this property. It can reactivate detailed traces of specific past experiences and has a substantially higher capacity than WM [6] . EM has traditionally been studied as a functionally and neurally separate system from WM [1, 6–9]. Accordingly, episodic contributions to WM tasks have often been treated as confounds to be controlled rather than mechanisms to be characterized [10]. Some prior work has shown that performance shifts from WM- to EM-driven once WM-capacity is exceeded, establishing that EM can supply otherwise-unavailable content to WM tasks [11]. However, it remains unknown which computations let EM contribute to WM discrimination in the first place, and when EM’s contribution actually occurs. EM and WM may further operate as an integrated system. For instance, spontaneous EM reinstatement during active maintenance has been shown to interfere with WM content even when unneeded [12], demonstrating interactions between EM and WM beyond on-demand recruitment. Understanding the integration between EM and WM thus requires a paradigm in which WM- and EM-representations are combined in a way that can identify which EM computations support WM, and when.

We propose that EM supports WM discrimination through three computations, each operating at a different time point. At encoding, pattern separation stores highly similar experiences as distinct traces allowing similar memory content to be discriminated [13–15]. Before a similar item reappears, predictive recall could then anticipate its content in advance. For instance, intracranial recordings during repeated naturalistic story-listening show spontaneous anticipatory neural activity on a second exposure [16] (see also [17]), suggesting EM can be recruited automatically and fast enough to resolve upcoming ambiguity. Finally, at the moment an item must be discriminated, pattern completion could recover its matching trace from a partial cue via autoassociative retrieval [18–21].

To probe these mechanisms, we developed the MST-back paradigm, combining mnemonic discrimination judgments of visually similar everyday objects [22] with a continuous N-back task [23]. Participants, undergoing eye-tracking, compared each object to its *n*^th^ predecessor, classifying it as “same”, “similar”, or “new”. A low-load one-back phase served as incidental encoding (Figure 1b); some pairs then reappeared in a subsequent high-load two-back phase (Figure 1c). We manipulated how pairs were experienced during encoding: *compared* pairs appeared consecutively (A1, B1), enabling direct comparison; *isolated* pairs were separated by at least five intervening items; *novel* pairs appeared for the first time at retrieval (A2, B2). This created a gradient for a potential EM benefit: *novel* items had no EM traces, *isolated* items could benefit from two disjoint EM traces, whereas *compared* items could leverage an EM of the contrasted experience. Importantly, item presentation during two-back was carefully balanced such that recognizing a previously encountered item could not predict the timing or type of the upcoming response. This ensured that EM could not alleviate processing demands by restricting which items needed to be maintained in WM.

**Fig. 1.**
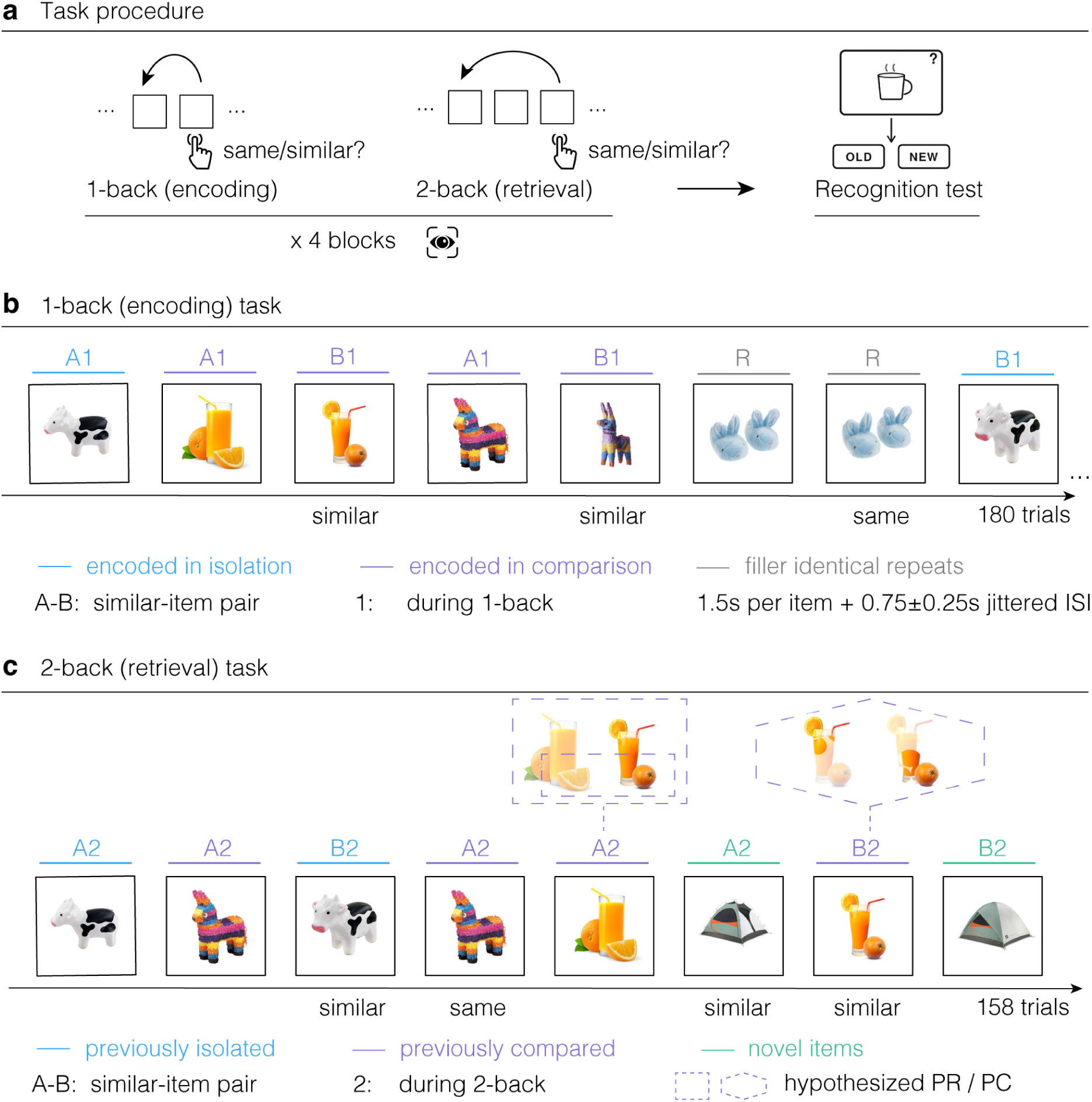
The MST-back task and experimental paradigm. **a** Task procedure: In two task phases, participants judged each item relative to its n*^th^* predecessor as “same”, “similar”, or “new” (no response). Each block contained a one-back phase (used to elicit episodic memory encoding) followed by a two-back phase, in which some items from the one-back phase reappeared to test for episodic retrieval effects. Participants completed four task blocks while undergoing eye tracking. A final old/new recognition test measured long-term memory for items encountered during one-back. Notation: A and B denote the two members of a perceptually similar pair, numbers (1 and 2) denote the task phase in which the item appeared (one-back vs. two-back). **b** One-back phase. In the *compared* condition, A1 and B1 appeared on consecutive trials, allowing direct comparison between similar items. In the *isolated* condition, A1 and B1 were separated by at least 5 intervening items, limiting direct comparison. Gray items indicate filler identical repeats (R). Each item was presented for 1.5 s followed by a jittered inter-stimulus interval of 0.75 plus or minus 0.25 s. **c** Two-back phase. Previously *compared*, previously *isolated*, and *novel* pairs appeared as A2 and B2, separated by an intervening item, within a continuous two-back stream. Thought bubbles above A2 and B2 indicate the hypothesized episodic processes supporting B2 discrimination: predictive recall (PR; rectangle), the anticipatory reinstatement of the upcoming pairmate at A2, and pattern completion (PC; hexagon), the reinstatement of the previously encountered item from a partial cue (here: partial sampling shown as a faded item with only the sampled regions at full opacity). Purple color indicates previously *compared* pairs, blue indicates pairs previously seen in *isolation*, green indicates *novel* pairs. Additional repeated trials were included such that recognizing A2 could not predict the timing or type of the upcoming response.

The three proposed computations predict a specific temporal profile of EM’s involvement in WM, and eye-tracking offers a trial-level readout to pinpoint each of them as they unfold. For instance, pupil dilation indexes the recognition of previously studied items [24] and the discrimination of similar lure items from previously studied targets [25], making it a candidate physiological correlate to characterize EM pattern completion. EM further guides eye movements toward memory-relevant locations [26]. Indeed, the overlap between an earlier viewing pattern and a later one acts as a marker of successful EM retrieval – a phenomenon known as gaze-pattern reinstatement [27, 28]. Gaze similarity may consequently track pattern completion in the MST-back, distinguishing successful from unsuccessful discrimination at response time. Moreover, trial-resolved *predictive* EM contributions may manifest in anticipatory viewing patterns: similarity between gaze on an item at retrieval and its similar pairmate’s earlier fixation pattern can be evaluated before that pairmate has reappeared, indexing predictive recall of upcoming content. In the one-back phase, the similarity between fixations on the two pairmates further offers a trial-level readout of how distinctly the pair was encoded.

### Episodic memory rescues working memory discrimination

Before testing for episodic contributions, we first confirmed that participants could perform the one-back and two-back tasks as intended. In the one-back phase, where each item was compared to its immediate predecessor under minimal WM load, participants performed with high accuracy across all required response types (*same*: mean = 89%, SD = 10.4%; *similar* : mean = 80%, SD = 13.1%; *new* : mean = 98%, SD = 2.2%; see Supplementary Figure S1 for details). Participants were further able to successfully perform the two-back task, providing accurate responses for 81% (SD = 16.6%) of repeated items (*same* response), 47% (SD = 11.9%) of similar items, and 99% (SD = 1.2%) of new items (see Supplementary Figure S2 for details).

We then examined, whether EM could rescue performance when WM demands were high. We predicted that, in the two-back phase, EM would support discrimination of similar items in WM, especially, if those items had been directly compared to each other in the one-back phase. We reasoned that this direct comparison in the one-back phase would promote the encoding of diagnostic item features. Indeed, the discrimination index – the difference between accuracy on *similar* trials and erroneous *similar* responses on *new* trials – was significantly modulated by encoding condition (Figure 2a; *F* (2, 60) = 46.99, *p < .*001, 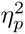 = .610, *ε* = .907, BF_10_ *>* 1000). Mnemonic discrimination was best for similar two-back items that had already been compared to each other in the one-back task, discrimination was at an intermediate level for similar items that had been presented in *isolation*, and it was *lowest* for *novel* items that appeared for the first time in the two-back task (*compared* vs. *isolated* : *t*(30) = 7.07, *d* = 1.27, *p < .*001; *isolated* vs. *novel* : *t*(30) = 3.59, *d* = 0.64, *p < .*001; *compared* vs. *novel* : *t*(30) = 8.53, *d* = 1.53, *p < .*001). Conversely, participants’ accuracy in detecting same item repetitions across the two-back lag (*d^′^*) did not differ between conditions (Figure 2b; *F* (2, 60) = 0.29, *p* = .750, 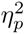 = .010, *ε* = .998). A Bayes factor analysis confirmed this, providing strong evidence for the null-hypothesis that having encountered an item in the one-back task did not increase participants’ ability to successfully identify item repetitions in the two-back phase (*same* responses; BF_01_ = 23.01). This null result indicates that the EM benefit was specific to discriminating between similar items, rather than reflecting a general improvement in memory for items encountered during one-back. This finding rules out accounts based on general familiarity.

**Fig. 2.**
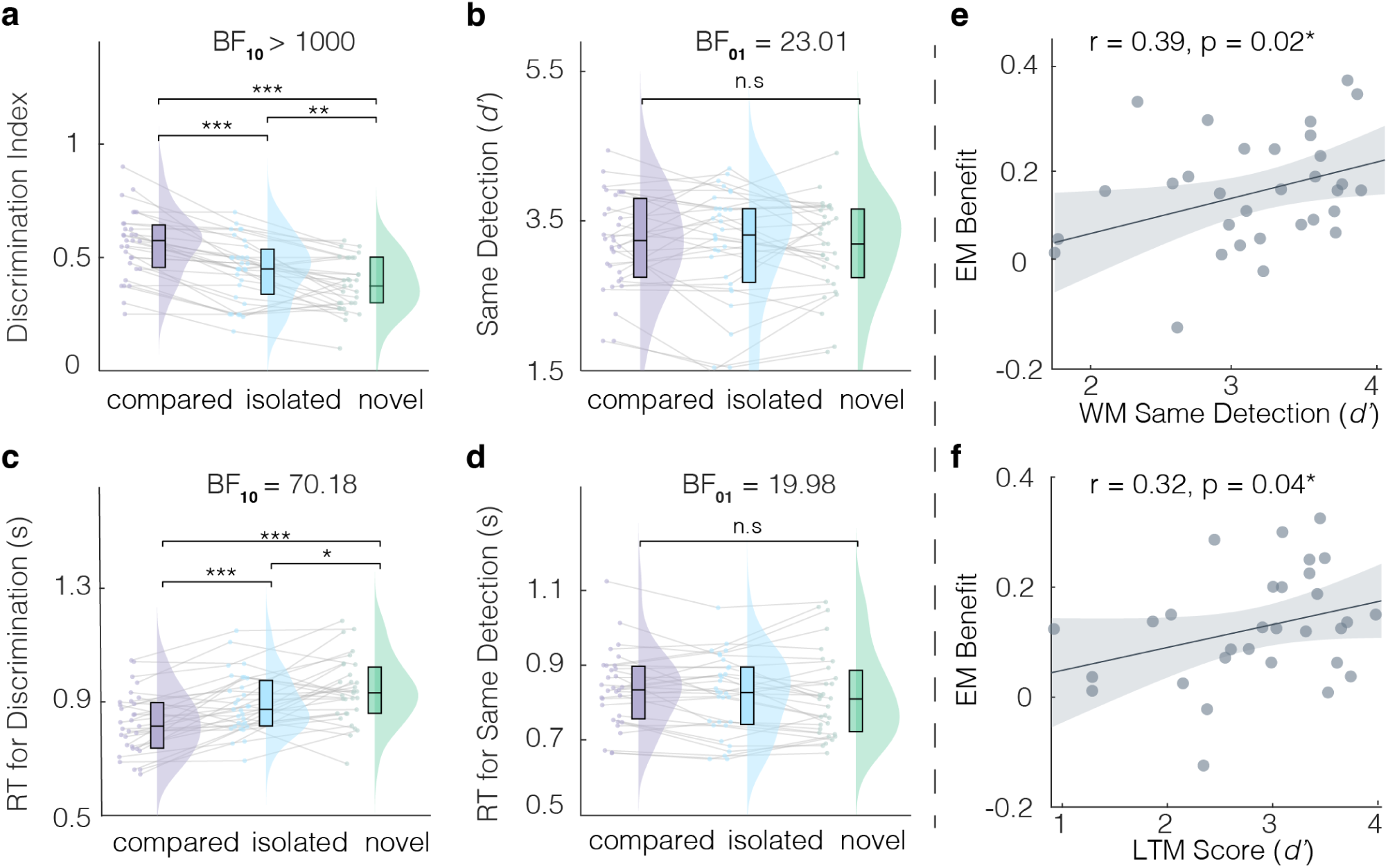
Episodic rescue of working memory discrimination is modulated by encoding condition. Behavioral performance during the two-back retrieval phase is shown as a function of encoding condition. In **a** through **d**, purple indicates items *compared* during one-back (EM formed in comparison), blue indicates items presented in *isolation* during one-back (separate episodic memories), green indicates *novel* items (WM only). **a**, Discrimination index for similar items. Performance was highest for *compared* items, intermediate for *isolated* items, and lowest for *novel* items. **b**, Same detection *d^′^*. Detection of identical repeats did not differ across encoding conditions. **c**, Median response time for correct similar item discrimination: responses were fastest for *compared* items, intermediate for *isolated* items, and slowest for *novel* items. **d**, Median response time for correct same item detection: response times did not differ across encoding conditions. **e, f**, The magnitude of the episodic benefit correlated across participants with both same-detection *d^′^* (e) and long-term recognition strength (f), consistent with a common factor linking WM and EM. Dots show individual participants, gray lines connect repeated measures, raincloud plots show group distribution. Box plots are median and interquartile range. Bayes factors quantify evidence for the alternative hypothesis (BF_10_) or for the null hypothesis (BF_01_). Asterisks indicate significance of comparisons (fdr-corrected).

Response times provided a complementary line of evidence. Correct mnemonic discrimination was fastest for *compared*, intermediate for *isolated*, and slowest for *novel* items (Figure 2c; *F* (2, 60) = 23.11, *p < .*001, 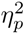 = .435, *ε* = .942, BF_10_ = 70.18; *compared* vs. *isolated* : *t*(30) = *−*4.61, *d* = 0.83, *p < .*001; *isolated* vs. *novel* : *t*(30) = *−*2.57, *d* = 0.46, *p* = .008; *compared* vs. *novel* : *t*(30) = *−*6.46, *d* = 1.16, *p < .*001), whereas response times for correctly identified same items did not differ between conditions (2d; *F* (2, 60) = 1.72, *p* = .192, 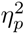 = .054, *ε* = .896, BF_01_ = 19.98), mirroring the same specificity seen in accuracy data. The convergence of faster and more accurate responses for compared items indicates that the EM benefit reflects genuine facilitation rather than a shift in response strategy.

Together, these results show that EM does more than modestly assist WM: when items had been directly compared during encoding, participants discriminated them more accurately and more quickly during high-load retrieval, and this benefit was absent for simple recognition of identical repeats. This pattern indicates that detailed episodic representations of similar items were integrated into the WM comparison process itself, rather than merely tagging items as previously encountered.

If the EM benefit reflects a general capacity for using memory to aid discrimination, its magnitude should track independent measures of both WM and long-term memory ability across individuals. We therefore asked whether the episodic benefit on similar trials scaled with participants’ WM and EM performance. To this end, we correlated the episodic benefit – estimated as the difference in mnemonic discrimination between old (*compared* and *isolated* condition) and *novel* items – with measures of pure WM and EM. Pure WM performance was estimated on same item detection in the two-back phase (WM-*d^′^*), since this measure was unaffected by encoding condition, making it a proxy for WM capacity uncontaminated by episodic support (see above). Long-term memory performance was derived from the final recognition task (EM-*d^′^*). Across participants, the magnitude of the episodic benefit significantly correlated with both, WM capacity (*r*(29) = .39, *p* = .02, Figure 2e), and long-term recognition accuracy (*r*(28) = .32, *p* = .04, Figure 2f). WM and long-term recognition performance were themselves strongly correlated (*r*(28) = .74, *p < .*001.) Together, these significant correlations indicate that the observed benefits scale with individual differences in working- and long-term memory performance consistent with a common factor contributing to WM, EM, and their interaction.

### Pupil dilation strength tracks episodic strength and successful discrimination

Next, we examined whether participants’ pupil dilation tracked the reinstatement of episodic memories during successful discrimination. We predicted that pupil dilation would scale with the strength of the episodic trace available. Thus, it would be greatest for compared items, intermediate for isolated items, and smallest for novel items. Further, we predicted an increase on trials where discrimination was successful, reflecting the recovery of useful information [26]. To test this, we compared the change in pupil size after stimulus onset between the *novel*, *isolated* and *compared* condition on trials that required a *similar* response. On these trials, the omnibus test revealed a single significant cluster extending from 0.951 to 1.499 s after stimulus onset (cluster-level *p <* 0.001, summed *F* = 9,912.08; see Analyses of pupil dilation and Supplementary Figure S3a) within which pupil dilation was significantly modified by encoding condition (Figure 3a). The three pairwise contrasts each yielded a significant cluster overlapping this interval: *compared* vs. *isolated*, 1.024–1.499 s (cluster-level *p* = 0.020, summed *t* = 1,596.08; *compared > isolated* ); *compared* vs. *novel*, 0.945–1.499 s (cluster-level *p* = 0.005, summed *t* = 2,953.77; compared *> novel* ); and *isolated* vs. *novel*, 1.002–1.499 s (cluster-level *p* = 0.018, summed *t* = 1,611.32; *isolated > novel* ). We summarized pupil dilation as the mean change from 1.0 to 1.5 s, bounded by the approximate cluster onset and the stimulus offset. A 3 *×* 2 follow-up ANOVA (encoding condition *×* discrimination accuracy) on the average pupil dilation within this 1.0–1.5 s window revealed significant main effects of encoding condition (*F* (2, 56) = 4.18, *p* = .023, 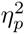 = .130, *ε* = .932, BF_10_ = 3.36) and accuracy (*F* (1, 28) = 40.58, *p < .*001, 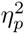 = .592, BF_10_ = 123.34), and a significant interaction (*F* (2, 56) = 10.78, *p < .*001, 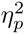 = .278, *ε* = .913). Consistent with our first prediction, pupil dilation was greatest for *compared* items, intermediate for *isolated*, and smallest for *novel* items (*compared* vs. *isolated* : *t*(28) = 3.56, *d* = 0.66, *p < .*001; *isolated* vs. *novel* : *t*(28) = 3.45, *d* = 0.64, *p < .*001; *compared* vs. *novel* : *t*(28) = 5.96, *d* = 1.11, *p < .*001).

**Fig. 3.**
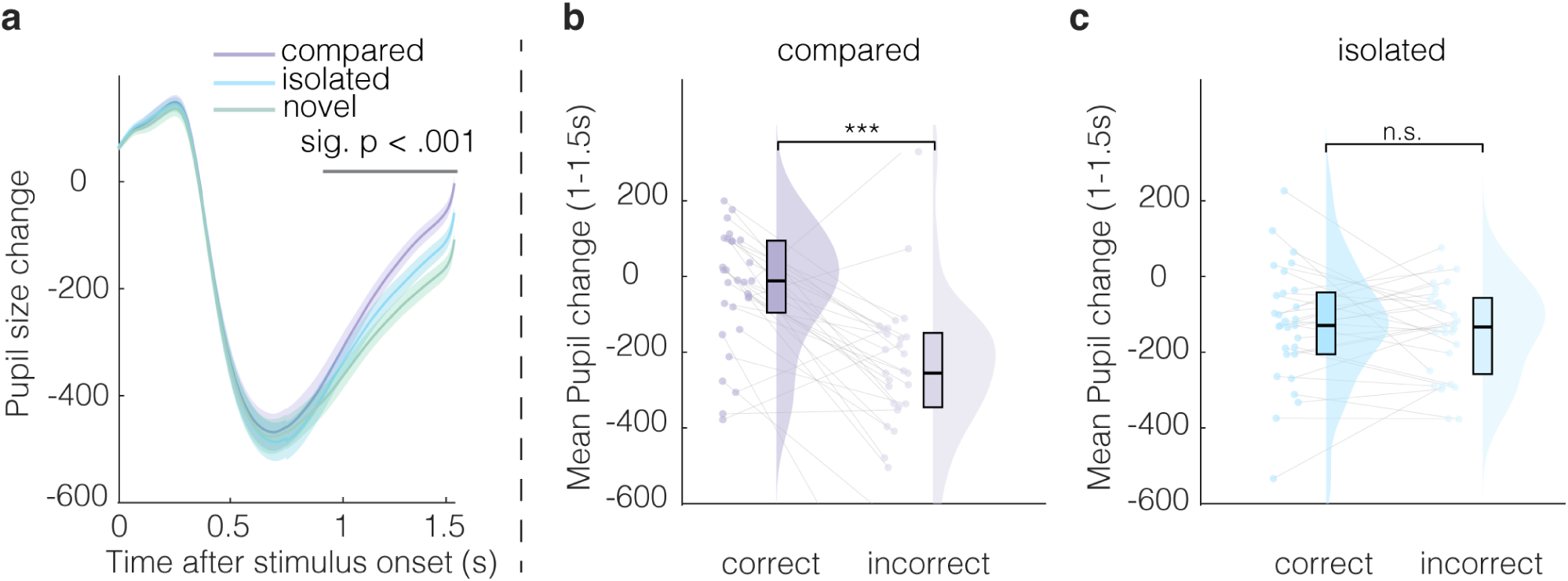
Pupil dilation marks episodic benefit during similar item discrimination. **a**, Change in pupil size after stimulus onset during similar item discrimination in the two-back phase. Lines show group averaged pupil change for *compared*, *isolated*, and *novel conditions*. Shaded regions show the standard error of the mean (*±* SEM across *n* = 29 participants). The gray line marks the cluster of significant modulation by condition. **b, c**, Mean pupil change between 1.0 and 1.5 s after stimulus onset for compared (**b**) and isolated (**c**) items, plotted separately for correct and incorrect discrimination trials. Successful discrimination was associated with larger pupil dilation in the compared condition, but not in the isolated condition. This dissociation indicates that the pupil response tracks the recruitment of a specifically comparative episodic trace, rather than memory strength in general. In b and c, dots show individual participants, gray lines connect participant values across accuracy bins, raincloud plots show group distribution, and box plots show the median and interquartile range.

Consistent with our second prediction, this gradient was accompanied by a further boost in dilation specifically on trials where discrimination succeeded: pupil dilation was greater on correct than incorrect trials in the *compared* condition (*t*(28) = 5.07, *d* = 0.94, *p < .*001) but this effect was absent in the *isolated* condition (*t*(28) = 0.62, *d* = 0.12, *p* = .27, BF_01_ = 4.24). Together, these results indicate that pupil dilation is not simply a marker of recognition in general, but tracks the successful recruitment of a sufficiently strong episodic trace — one formed through direct comparison at encoding.

To test whether this modulation was specific to discrimination rather than a general effect of encoding condition on pupil response, we repeated the same analysis on trials requiring a “same” response. On these trials, no cluster exceeded the cluster-forming threshold for the omnibus effect or for any pairwise contrast; the 1.0–1.5 s window derived from discrimination trials was therefore applied unchanged to the same-detection control analysis; A one-way repeated-measures ANOVA on the average pupil dilation within the same 1.0–1.5 s window revealed no effect of condition (Supplementary Figure S3c; *F* (2, 56) = 1.00, *p* = .373, 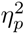 = .034, *ε* = .959, BF_01_ = 19.26). This null result indicates that the pupil effects above were specific to trials requiring mnemonic discrimination, and cannot be explained by generic differences in arousal, effort, or task engagement across encoding conditions.

### Gaze patterns carry image-specific information

We next examined participants’ trial-by-trial fixation patterns (gaze patterns) to characterize the mechanisms underlying EM contributions to task performance. Before testing the relationship between gaze patterns and performance, we first confirmed that gaze patterns carried sufficient item-specific information to distinguish individual images from their similar pairmates. We verified that gaze similarity between different viewings of the same item (A1–A2 and B1–B2 similarity), and between different viewings of similar-but-different items (A1– B1, A1–B2, A2–B1, A2–B2) exceeded a permutation-based null distribution computed from different viewings of unrelated items (Figure 4; all *p < .*001). This confirms that fixation patterns carry item-specific representational content at sufficient resolution during image viewing.

**Fig. 4.**
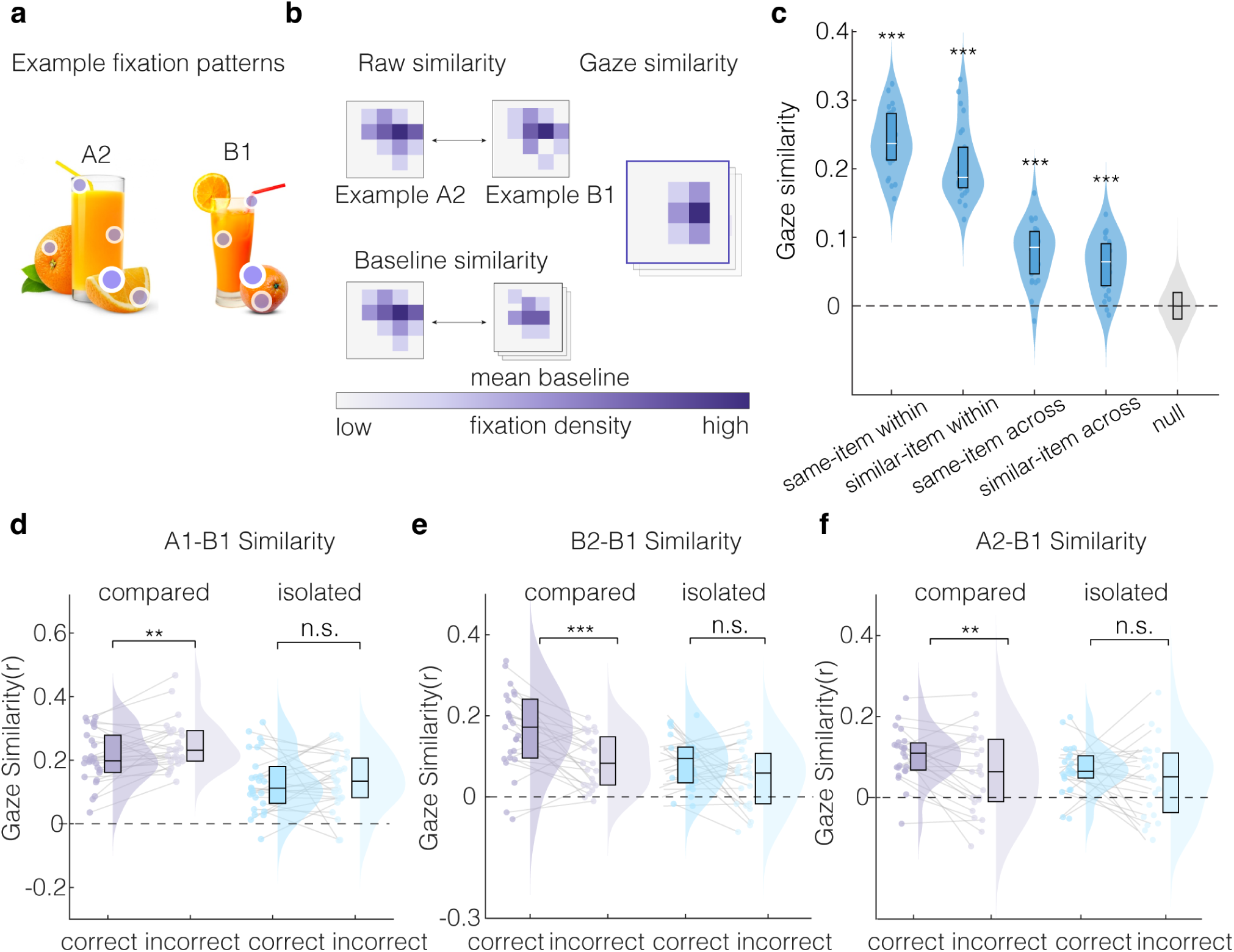
Gaze similarity tracks episodic contributions to working memory. **a**, Example fixation patterns from the viewing of two similar pairmates during A2 (left) and B1 (right) trials. Purple dots with white outlines indicate real fixations. Longer fixations within a region are illustrated as darker color. **b**, Gaze similarity computation. Fixation density maps were generated for each trial. Raw gaze similarity was computed as the Pearson correlation between two fixation density maps (e.g., A2 to B1). Baseline similarity was computed as the average correlation between trials with unrelated items, within a given comparison (e.g., A2 to unrelated B1), to account for unspecific viewing patterns. Baseline-corrected gaze similarity was obtained by subtracting this baseline from raw similarity between trial-combinations of interest. The heatmaps illustrate fixation density and corrected similarity; the color scale indicates relative fixation density. **c**, Baseline-corrected gaze similarity across item relationships. Same item comparisons within phases (A1 to A1, A2 to A2), similar item comparisons within phases (A1 to B1 and A2 to B2), same item comparisons across phases (A1 to A2 and B1 to B2) similar item comparisons across phases (A1 to B2 and A2 to B1) – shown left to right as plotted – all exceeded the permutation based null distribution (centered at zero, as per baseline correction procedure). This indicates that fixation patterns contained item-specific and relationship-specific information. **d–f**, Note that lower similarity in d indicates more distinct encoding of the two pairmates, whereas higher similarity in e and f indicates stronger reinstatement of an earlier fixation pattern **d**, A1 to B1 gaze similarity indexes differential processing of two similar items during encoding (pattern separation). In the *compared* condition, A1 to B1 similarity was lower for pairs that were later discriminated correctly. No comparable subsequent memory effect appeared in the *isolated* condition. **e**, B2 to B1 gaze similarity indexes reinstatement of B’s one-back fixation pattern when B is encountered during two-back. In the *compared* condition, B2 to B1 similarity was greater on correct than incorrect discrimination trials, indicating pattern completion of discriminative viewing patterns. No comparable effect appeared in the *isolated* condition. **f**, A2 to B1 gaze similarity indexes reinstatement of the counterpart’s (B item) one-back fixation patterns during the presentation of A in the two-back task. In the *compared* condition, A2 to B1 similarity was greater on trials where the corresponding B2 discrimination was successful, compared to trials with incorrect later B2 discrimination, suggesting predictive recall of upcoming information. No comparable effect appeared in the *isolated* condition. In **d** through **f**, dots show individual participants, gray lines connect participant values, raincloud plots show distributions, and box plots are medians and interquartile ranges.

### Gaze pattern separation at encoding predicts subsequent discrimination success

In order to enable EM to support discrimination, representations of the differences between images would need to be laid down at encoding. We therefore examined how participants’ gaze similarity between the A and B items of a similar pairmate differed between trials that were successfully discriminated in the subsequent two-back trials, and those trials that were not classified correctly (i.e. erroneous same responses and omissions). We hypothesized that encoding items in the *compared* condition would produce more distinctive sampling of the pairmates than encoding them separately, since direct comparison allows diagnostic features to become salient during viewing. This is the gaze-based signature of pattern separation: lower similarity between fixations on the two items should reflect more distinct encoding, and predict better subsequent discrimination. To test this, we examined the difference in gaze similarity between the A and B items of a pair between trials that were subsequently discriminated correctly in the two-back phase, and those that were not (i.e., erroneous same responses and omissions).

A 2*×*2 ANOVA (encoding condition *×* subsequent B2 accuracy) on A1–B1 gaze similarity revealed a significant main effect of encoding condition (*F* (1, 25) = 33.65, *p < .*001, 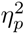 = .57), with no main effect of accuracy (*F* (1, 25) = 1.83, *p* = .188, 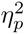 = .07) and no interaction (*F* (1, 25) = 2.48, *p* = .128, 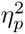 = .09). Planned simple effects on subsequent accuracy revealed that, in the *compared* condition, A1–B1 gaze similarity was significantly lower, indicating distinct encoding of the two items, on trials where B2 was subsequently correctly discriminated (Figure 4d; *t*(25) = *−*2.53, *d* = *−*0.50, *p* = .009); no such effect emerged in the isolated condition (*t*(25) = *−*0.19, *d* = *−*0.04, *p* = .427, BF_01_ = 4.15). These results suggest that participants who sampled A1 and B1 more distinctively during encoding produced more separable memory representations, supporting later discrimination. Note that an alternative account — in which intrinsically distinctive pairs simply yield both lower gaze similarity and higher accuracy — is unlikely to explain this pattern: pair-to-condition assignment was randomized across participants, so any such item-driven effect should appear in both conditions rather than only the compared condition.

### Gaze pattern completion at the moment of discrimination

We next examined whether item-specific episodic memories were reactivated in gaze patterns at the moment of the *similar* discrimination response (B2 item). We predicted that item B’s one-back representation would be reinstated during its two-back perception, guiding gaze toward the image’s diagnostic features. A 2 *×* 2 ANOVA (encoding condition *×* discrimination accuracy) on B2–B1 gaze similarity revealed significant main effects of encoding condition (*F* (1, 23) = 24.50, *p < .*001, 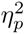 = .52) and accuracy (*F* (1, 23) = 13.63, *p* = .001, 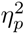 = .37), with no interaction (*F* (1, 23) = 1.20, *p* = .285, 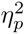 = .05)). B2–B1 gaze similarity was greater in the compared than the isolated condition (*t*(23) = 5.69, *d* = 1.16, *p < .*001), and was greater on correct than incorrect trials in the *compared* condition (Figure 4e; *t*(23) = 3.63, *d* = 0.74, *p < .*001); the same contrast in the *isolated* condition did not reach significance, although a statistical trend was observed (*t*(23) = 1.71, *d* = 0.35, *p* = .051). This partial trend in the isolated condition stands in contrast to the separation and predictive-recall effects reported above (Figure 4d, f), which were absent in the isolated condition entirely, suggesting that pattern completion, unlike the other two mechanisms, may be only partially dependent on direct comparison at encoding. These results confirm that the reinstatement of gaze patterns characteristic of the B item during one-back task marked its correct discrimination during two-back; this was statistically significant in the *compared* condition.

### Predictive recall of gaze patterns marks upcoming discrimination success

Finally, we hypothesized that successful discrimination of a similar item would be supported by predictive recall of the upcoming similar B2 item during presentation of the A2 item in the two-back task. Because EM encoding in the one-back task makes the distinguishing features of an item pair available, we expected that participants would sample the A item during two-back in a way that anticipates key features of the B item. Hence, we modeled accuracy – determined by the response to the counterpart (B2) – based on the A2–B1 similarity of gaze patterns.

As expected, a 2 *×* 2 ANOVA (encoding condition *×* discrimination accuracy of the B2 response) on A2–B1 gaze similarity revealed a significant main effect of accuracy (*F* (1, 22) = 5.03, *p* = .035, 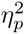 = .19), with no main effect of condition (*F* (1, 22) = 1.66, *p* = .210, 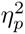 = .07) and no interaction (*F* (1, 22) = 0.59, *p* = .452, 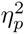 = .03)). Planned simple effects revealed that A2–B1 gaze similarity was significantly greater on correct than incorrect trials in the *compared* condition (Figure 4f; *t*(22) = 2.59, *d* = 0.54, *p* = .008), whereas no comparable effect emerged in the *isolated* condition (*t*(22) = 0.84, *d* = 0.17, *p* = .206).

In order to characterize how predictive recall at A2 unfolds over time, we interrogated A2–B1 similarity at cumulative fixations. To this end, we computed cumulative gaze similarity across successive fixations at A2 and estimated the slope of accumulation, thereby providing a readout of how the representation of the episodic memories incrementally emerges during predictive recall (Figure 5a).

**Fig. 5.**
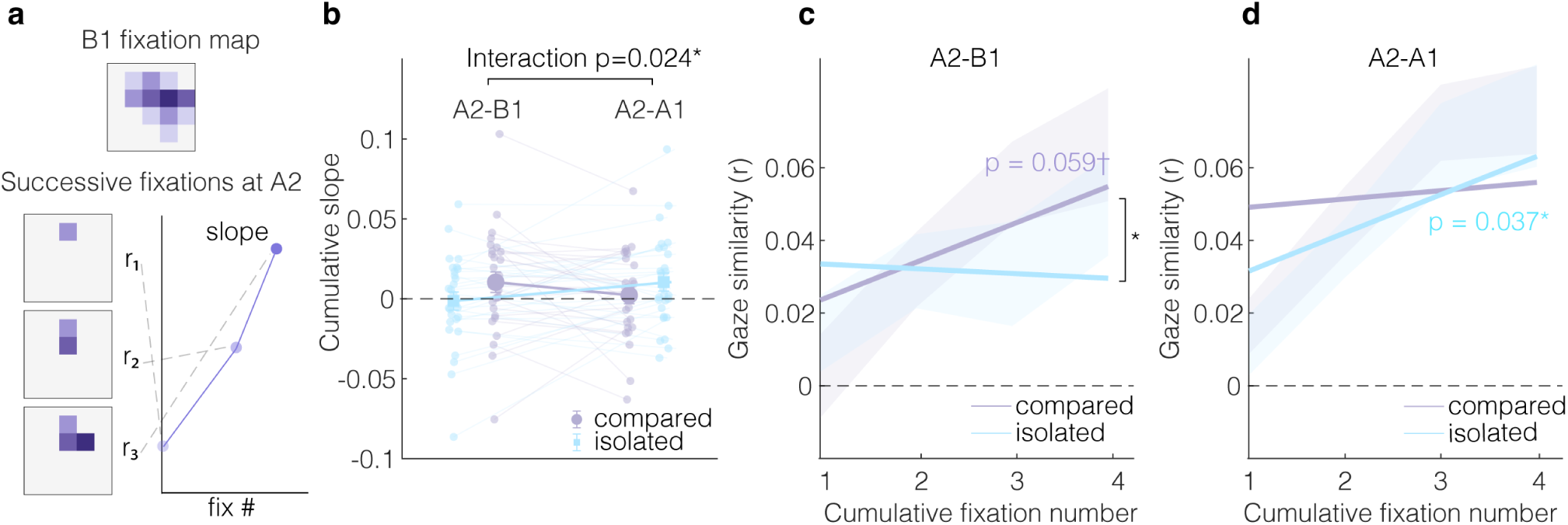
Predictive recall of compared pairmates emerges gradually but isolated items are recapitulated. Purple indicates the compared condition and blue indicates the isolated condition throughout. **a**, Schematic of the cumulative gaze similarity analysis. For each A2 retrieval trial, fixation density maps were constructed cumulatively across successive fixations at A2 in the two-back task. Cumulative maps from fixations 1 to 4 (bottom) were correlated with the corresponding one-back fixation map (top) for each similarity measure (B1 for A2–B1 similarity; A1 for A2–A1 similarity) and then baseline-corrected. **b**, Participant-level cumulative slopes for A2–B1 and A2–A1 gaze similarity. Slopes were estimated as the regression of gaze similarity over cumulative fixation number within each trial and then averaged within participant and condition. A 2 by 2 ANOVA on cumulative slopes showed a significant interaction between encoding condition and similarity type. This interaction indicates that compared encoding selectively increased cumulative reinstatement of the counterpart representation, A2 to B1, while isolated encoding preferentially supported accumulation of the item’s own representation, A2 to A1. Dots show individual participants, gray lines connect participant values across conditions. **c**, Cumulative A2 to B1 gaze similarity, indexing predictive recall of the perceptually similar item’s one-back gaze-pattern. In the *compared* condition, A2 to B1 similarity accumulated across fixations at trend level (in-panel *p*), whereas the *isolated* condition showed no reliable accumulation. Accumulation was significantly greater in the *compared* than *isolated* condition (bracket). **d**, Cumulative A2 to A1 gaze similarity, indexing recapitulation of the current item’s one-back fixation pattern. The *isolated* condition showed significant positive accumulation, whereas the *compared* condition did not, and the two conditions did not differ. In **c** and **d**, lines show group means and shaded bands denote 95% confidence intervals. Symbols: *†* indicates *p <* .10, * indicates *p <* .05, tested against zero for in-line annotations and compared between conditions for bracketed contrasts.

Predictive recall would need to emerge over the course of the trial, since retrieval demand is determined by perception of the A2 item. We further predicted that this gradual emergence would take a different form depending on encoding history: in the *compared* condition, where A2–B1 similarity distinguished successful from unsuccessful discrimination, gaze at A2 should increasingly resemble the not-yet-reappeared B item. In the *isolated* condition, where no such pairmate-anticipation effect was observed, gaze at A2 should instead show gradually emerging similarity to the item’s own early (A1) fixation pattern, reflecting completion of its own trace rather than anticipation of its pairmate.

A 2 *×* 2 ANOVA on cumulative slopes (pair type [A2–B1, A2–A1] *×* encoding condition [compared, isolated]) yielded a significant interaction (Figure 5b; *F* (1, 26) = 5.78, *p* = .024, 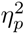 = .182), indicating that compared and isolated encoding produced distinct cumulative reinstatement profiles; no main effect of pair type (*F* (1, 26) = 0.20, *p* = .655, 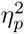 = .008) or encoding condition (*F* (1, 26) = 0.12, *p* = .731, 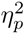 = .005) was observed. This result confirmed that gaze-similarity to the not-yet-reappeared B item and to the item’s own early (A1) fixation pattern accumulated differently in the *compared* and *isolated* condition. In line with our first prediction, we found a statistical trend indicating that A2–B1 similarity accumulated across fixations in the *compared* condition, while the *isolated* condition did not provide statistical evidence for accumulation of A2–B1 similarity (Figure 5c; compared: *M* = 0.010, *SD* = 0.034, *t*(26) = 1.62, *p* = .059; isolated: *M* = *−*0.001, *SD* = 0.030, *t*(26) = *−*0.23, *p* = .591). In direct comparison, evidence for accumulation of A2–B1 similarity was significantly greater in the *compared* condition than in the *isolated* condition (*t*(26) = 2.34, *d* = 0.45, *p* = .014), lending further support to the interpretation that gradually emerging A2–B1 similarity was unique to the *compared* condition. Conversely, A2–A1 similarity followed a different pattern: it accumulated in the *isolated* condition but it did not significantly accumulate in the *compared* condition (Figure 5d; *compared* : *M* = 0.002, *SD* = 0.028, *t*(26) = 0.43, *p* = .336; *isolated* : *M* = 0.011, *SD* = 0.029, *t*(26) = 1.86, *p* = .037), supporting our second prediction that items encoded in isolation would be characterized by gradual pattern completion of their own trace rather than pairmate-anticipation. A direct comparison of the conditions, however, did not yield evidence for a significant difference in accumulation of A2–A1 similarity (*compared* vs. *isolated* : *t*(26) = *−*1.07, *d* = *−*0.21, *p* = .852). Taken together, these results suggest that gaze patterns characteristic of the anticipated B item emerge gradually when a memory of their comparison has been formed. Conversely, when both items have been encoded in isolation, gaze patterns characteristic of the original experience are recapitulated.

## Discussion

We asked whether, and how, episodic memory (EM) can rescue working memory (WM) discrimination in continuous experience. Using the MST-back paradigm – a stream of object images in which participants detect recurrences of identical or similar items – we manipulated memory traces during a one-back encoding phase by presenting similar item pairs either in isolation or in direct comparison. We then isolated the contribution of EM to WM by contrasting two-back mnemonic discrimination for previously *compared*, previously *isolated*, and *novel* items. Discrimination of similar items was best for pairmates in the *compared* condition; encoding in *isolation* produced a smaller but still significant benefit over *novel* pairs. Crucially, the benefit of EM was specific to the discrimination of similar items; we found no such benefit on trials requiring the identification of identical repeats. This benefit was further accompanied by greater pupil dilation when discrimination was successful in the *compared* condition. Moreover, we identified three markers of EM in participants’ gaze patterns: During one-back encoding, A1-B1 similarity was reduced for those pairmates that were later discriminated correctly (pattern separation), while during two-back retrieval, greater B2-B1 gaze similarity indicated reinstatement at the moment of successful discrimination (pattern completion) and anticipatory reinstatement of the not-yet-visible pairmate (greater A2-B1 similarity) predicted successful discrimination two trials in advance (predictive recall).

Our central behavioral finding is that WM discrimination was significantly enhanced when prior encoding established an episodic representation that could supply diagnostic information, showing that episodic representations can be directly integrated with WM to rescue performance once capacity is exceeded. Contributions of EM to WM have previously been characterized at the level of aggregate task performance (e.g., [11, 29]); our behavioral findings go beyond these demonstrations by showing that this benefit appeared only on trials requiring mnemonic discrimination of similar items and was markedly absent on trials requiring detection of identical repeats — ruling out generic processing benefits from consecutive presentation, such as priming or sustained attention [30, 31]. We can also exclude accounts based on EM freeing up resources, since our trial structure was balanced such that item novelty was not predictive of the type or timing of the required response. Pupil dilation provided converging, independent evidence that EM contributed to WM discrimination. Dilation followed the same encoding gradient as behavioral performance — greatest for compared items, intermediate for isolated, and smallest for novel — and was larger on correct than incorrect trials in the compared condition. This pattern is difficult to explain by effort alone, since effort accounts typically predict greater dilation for harder or incorrect decisions [32, 33] — the opposite of what we observed. Instead, dilation appears to index the successful recovery of diagnostic episodic detail, consistent with prior work linking pupil dilation to EM engagement and retrieval [24, 34–36].

Examining gaze patterns further allowed us to track EM’s contribution to WM at a high level of temporal precision. Because fixations are spatially and temporally resolved, we were able to pinpoint the three computations that realized the episodic benefit at the moments they exerted their effects — pattern separation, pattern completion, and predictive recall: distinctive encoding of a pair predicted later discrimination success, reinstatement of an item’s own encoding pattern marked successful retrieval, and anticipatory reinstatement of the not-yet-visible pairmate preceded successful discrimination by two trials.

The staged episodic benefit on WM performance carries a broader implication: EM’s contribution to WM is not a single, undifferentiated boost, but an unfolding process in which distinct computations are recruited at the specific moments they are needed. Separation must occur at encoding to lay down a distinguishing trace; anticipatory recall can only precede an item when predictive information is available in memory; and pattern completion operates once retrieval is actually demanded. This intricate dependency reframes the relationship between the two systems: rather than EM acting as a supplementary resource that can step in as a backup, our results suggest that its computations are woven into WM processing at the timescale of individual comparisons, in a way that may be invisible to methods that assess performance only at the level of the block or the trial’s outcome.

This staged process was not, however, equally present across encoding conditions. All three gaze markers — along with the pupil dilation effect reported above — were confined to items that had been directly compared during encoding, even though isolated encoding still produced a smaller but reliable behavioral benefit over novel items. This asymmetry may reflect transfer-appropriate processing [37]: encoding the differences between two similar items head-to-head produces a trace whose format directly supports the later comparison, allowing it to guide gaze at retrieval, whereas isolated encoding produces two independent traces that may support recognition without leaving as clear a signature in viewing behavior. This is consistent with the hippocampal differentiation literature, where directly contrasting similar items pushes their representations apart and yields later advantages in discrimination [15, 38, 39].

Our findings also speak to the debate over activity-silent WM. WM content has been shown to persist across delays during which neural activity returns to baseline [40–42], a pattern difficult to reconcile with strict active-maintenance accounts [43]. The activity-silent framework attributes this persistence to transient synaptic modifications within WM itself [44], but persistent memory without active maintenance is equally consistent with an EM contribution [45], and the two accounts are difficult to distinguish from behavior alone. Our results suggest one way forward: because we tracked the reinstatement of episodic content directly, via gaze and pupillary markers tied to hippocampal computations [14, 17, 26], we could identify when EM was contributing to performance rather than inferring it from the persistence of behavior across a delay. This suggests that at least some instances of WM-like persistence attributed to activity-silent mechanisms may instead, or additionally, reflect EM contributions, a possibility that could be tested directly by applying a similar approach to tasks that have motivated activity-silent accounts.

Taken together, our findings support a novel perspective on working memory and episodic memory as a tightly integrated memory-system. They demonstrate that working memory, rather than operating as an isolated, capacity-limited system, may routinely draw on the same computations long recognized as central to episodic memory. Decades of treating WM and EM as separate objects of inquiry may have obscured a shared computational substrate that operates across both. Reuniting them, at both a theoretical and paradigmatic level, may be necessary to understand either system on its own terms, and human memory more broadly.

## Methods

### Participants

Thirty-one healthy adults (mean age = 20; range = 18-26; 19 female; all right-handed, with normal or corrected-to-normal vision) were recruited from the New York University Psychology Subject Pool. All participants provided informed consent electronically via Qualtrics upon arrival, and all procedures were approved by the New York University Institutional Review Board (IRB-FY2025-10338). Participants received course credit for their participation.

### Stimulus material

Stimuli were images of everyday objects drawn from the Mnemonic Similarity Task stimulus set [22], which provides pairs of perceptually similar object exemplars (denoted A and B throughout this manuscript based on their order of appearance). Each pair (1,192 pairs in total) consists of two exemplars of the same object (e.g., two glasses of orange juice) but differ in fine-grained perceptual features such as orientation, color, or local detail. Similarity ratings are available for all stimuli; these were obtained in the original study [22] from empirical confusion rates between A and B in a recognition test. To maximize discrimination difficulty and isolate fine-grained mnemonic processing, we selected 360 pairs from the two most perceptually similar stimulus groups and balanced pairs across these groups. All images were resized to 400 x 400 pixels, presented on a uniform gray background, and matched for mean luminance and root-mean-square contrast to control for low-level visual confounds.

### Apparatus and eye tracking

Stimuli were presented on a 27-inch LCD monitor (1920 × 1080 resolution, 60 Hz refresh rate) at a viewing distance of 110 cm. Each 400 × 400 pixel stimulus subtended approximately 8.0° × 8.0° of visual angle, presented centrally within a display that spanned approximately 30.6° × 17.4° of visual angle. Participants’ heads were stabilized using a chinrest with forehead support, and they were seated such that their eyes aligned with the upper quarter of the monitor.

Eye movements and pupil diameter were recorded monocularly from the right eye at 1000 Hz using an EyeLink 1000 desktop-mounted eye tracker (SR Research, Mississauga, Ontario, Canada). The eye tracker was positioned in accordance with the manufacturer’s recommended setup geometry at 50 to 55 cm from the participant’s chinrest, with its top knob centered horizontally on the front of the monitor and its height adjusted to maximize coverage without occluding the display.

The eye tracker was calibrated at the start of each task phase using a 9-point grid spanning the display (seven calibrations in total), with each calibration validated by a subsequent 9-point validation procedure. Calibrations were accepted only when mean spatial error fell below 1° of visual angle and maximum error below 1.5°. Otherwise, calibration was repeated. Drift was assessed at the start of each trial via a central fixation cross, and the eye tracker was recalibrated mid-block if drift exceeded 1° of visual angle.

The experiment was implemented in MATLAB using Psychtoolbox-3 [46–48], with stimulus timing synchronized to the eye-tracking recording via the EyeLink Toolbox [49]. Manual responses were collected via a standard keyboard.

### Overview of task procedure

The experiment consisted of four blocks of one-back and two-back phases, followed by a post-experiment old/new recognition test (Figure 1a). Within each block, participants first completed a continuous one-back mnemonic similarity task, followed by a 45-second inter-phase interval, and then a continuous two-back mnemonic similarity task. The two phases shared stimuli presentations, response options, and trial timing parameters; they differed only in the lag (*n*) at which each item was compared to its predecessor (*n* = 1 during one-back, *n* = 2 during two-back). This structural equivalence allowed episodic encoding and retrieval to emerge from the task demands themselves, without explicit study-test instructions that could differentially engage strategic processing across phases. Eye movements and pupil diameter were recorded continuously throughout all phases.

### Trial structure and timing

On each trial, a single image was presented centrally for 1.5 s, followed by a fixation cross for a jittered inter-stimulus interval of 0.75 ± 0.25 s (uniformly distributed). Participants classified the current item relative to its *n^th^*predecessor as “same” (identical repeat of the n-back item), “similar” (perceptually similar exemplar of the same object), or “new” (an item unrelated to the n-back item). Responses were entered by pressing “j” for same and “k” for similar; new responses required no keypress within the trial window (note that any keypress for new responses was identified as incorrect). Each block contained equal numbers of “same” and “similar” trials, equating j- and k-presses across conditions and participants. Because the key prompt disappeared throughout the actual trials, participants rested their right index and middle fingers on the “j” and “k” keys, respectively, for the duration of the task, and were instructed to memorize the key mappings before the task began, minimizing eye and head movements.

### One-back task

Each one-back phase consisted of 180 trials across four blocks (720 trials in total; Figure 1b). Participants classified each item relative to the immediately preceding item. The key manipulation controlled how members of each similar-item pair (A and B) were experienced. In the *compared* condition, A and B appeared on consecutive trials, such that the appearance of B required a “similar” response relative to A and afforded immediate comparison between the two exemplars. In the *isolated* condition, A and B were separated by at least five intervening unrelated items (median 50, range 5–178 across all participants), limiting the opportunity for direct comparison in WM; each item required its own classification relative to the immediately preceding item, which was always unrelated. Each block contained 30 compared pairs (60 trials; requiring 30 “similar” responses on the second presentation) and 30 isolated pairs (60 trials; both members required a “new” response, i.e., no keypress). To balance response frequencies and provide same-item detection events, additional 30 repeat pairs (60 trials) were embedded in which a single item appeared on two consecutive trials (requiring 30 “same” responses on the second presentation); these repeat items were unique to the one-back phase and did not appear in the subsequent two-back phase or in any A-B pair. Of the 180 trials per block, 30 required a “similar” response, 30 required a “same” response, and the remaining 120 required no response (“new”). Across the four blocks, the one-back phase contained 240 A-B pairs in total (120 compared, 120 isolated).

### Two-back task

Each two-back phase consisted of 158 trials across four blocks (632 trials in total; Figure 1c). Participants classified each item relative to the item presented two trials earlier (two-back). Similar-item pairs from one-back were tested in the two-back as A-X-A/B/N triplets, in which A appeared first, followed by an unrelated intervening item X, and then A, B, or N two trials later, where N was a different A. Participants were instructed to compare across the two-item lag (for example, juice A - backpack X - juice B, requiring a “similar” response to B). The retrieval phase introduced a third condition. Pairs in the *novel* condition consisted of items not previously encountered during one-back, presented for the first time in the two-back as A-X-A/B/N triplets. This structure matched the compared and isolated conditions, ensuring that item novelty alone could not predict whether the upcoming response required an “old”, “similar”, or “new” judgment. The two-back phase therefore tested three conditions: previously *compared* items (A and B encountered consecutively at encoding), previously *isolated* items (A and B encountered with at least five intervening items at encoding), and *novel* items (A and B encountered for the first time at retrieval). Across the four blocks, the two-back phase tested 360 A-B pairs in total, consisting of 120 previously *compared*, 120 previously *isolated*, and 120 *novel* pairs.

### Two-back trial structure and counterbalancing

Trial order in the two-back task was constructed to satisfy three constraints simultaneously. First, response requirements were balanced across conditions: of the 120 pairs per condition, 40 required a similar response (A-X-B, where B was the perceptually similar counterpart of A), 40 required a same response (A-X-A, where the second item was an identical repeat of the first), and 40 required no response (A-X-N, where the second item was unrelated to A). As a consequence, recognizing a previously encountered A item could not predict the upcoming response: a participant encountering juice A at retrieval was equally likely to encounter juice A again (same), a similar juice B (similar), or an unrelated object (new) two trials later.

Second, intervening items (X) within any triplet could again serve as the A item of another triplet, with their corresponding B appearing two trials later. Because any item in the retrieval stream could be the start of a new triplet whose B was forthcoming, participants could not free up WM resources by ignoring the middle item of a triplet, even when the first item of that triplet was recognized as previously encountered.

Third, the position of an item within a triplet (A vs. X vs. B) was unpredictable from any property of the item itself, including its encoding history (*compared*, *isolated*, *novel* ) and the response it would eventually demand. Whether a previously encountered item served as A in a new triplet or as X in another triplet was determined randomly, subject to the response-balancing constraint above. Because the two-back sequence contained *novel* items, this property further ensured that recognizing A items did not free up WM resources from the previous item, because a predicted intervening item X could reveal itself as the B item of the previous item. For each participant, pair assignment to encoding condition (*compared*, *isolated*, *novel* ), assignment of A or B as the encoded exemplar (relevant for the recognition test, see below), and trial order were independently randomized, subject to the constraints above.

Each block contained 30 triplets per condition, 90 in total. Because each intervening (X) item was itself the A item of another triplet, and each A-X-N triplet’s third (N) item initiated the subsequent triplet, these 90 triplets were realized in 150 stimulus trials: 90 first presentations of an A item, each requiring a *new* response, together with 30 *similar* (A-X-B) and 30 *same* (A-X-A) targets. With two filler items at the start of each block, five at its end, and occasional additional filler, each block comprised 158 trials. This additional filler was inserted near a block’s end, when a pending target required placement at the two-back lag but no unstarted triplet remained to occupy the intervening position.

In sum, this interleaved structure ensured that all items in the two-back stream remained relevant for discrimination, ensuring continuous WM engagement throughout the phase.

### Task instructions and practice

Before each phase was first encountered, participants completed on-screen instructions describing the task structure, response mappings, and trial timing. Instructions were followed by a practice block of 20 trials that matched in structure to the upcoming phase but used different stimuli from the ones employed in the main experiment. Participants were required to achieve at least 80% accuracy across all trials of the practice block to proceed to the main task; if accuracy fell below this threshold, the practice block was repeated until the criterion was met. Participants were given the opportunity to ask questions during instructions and between practice and main blocks; the experimenter provided clarification as needed.

### Recognition task

Following the four one-back/two-back blocks, participants completed a post-experiment old/new recognition test that provided an independent, long-term measure of memory for the encoded items. On each trial a single image was presented, and participants made a two-alternative forced-choice recognition judgment (“old” or “new”), indicating whether they had encountered it earlier in the experiment by pressing “j” for “old” or “k” for “new”. Each trial began with a 0.5 s central fixation, followed by the image, which remained on screen until a response was made, up to a maximum of 4 s. The old items were obtained by randomly selecting the A or B exemplar from each of 160 pairs – 80 from the *compared* and 80 from the *isolated* condition – that had been encountered during the one-back phase and whose similar pairmate (A-X-B) or identical repeat (A-X-A) reappeared during the two-back phase. These were intermixed with 160 novel foil images that had not appeared at any earlier point in the experiment. Because encoding during the one-back phase was incidental (see One-back task), this test was not announced in advance.

### Data exclusion

All exclusion criteria were defined before statistical analyses were conducted. Exclusion criteria were applied independently for behavioral, pupillometry, and fixation data.

#### Behavioral data exclusion

All participants were retained for behavioral analyses (N = 31), except for the final recognition test, which one participant did not complete (N = 30 for recognition analyses). Trials with reaction times shorter than 150 ms were excluded from reaction-time analyses as anticipatory responses were unlikely to reflect deliberate discrimination. Accuracy and reaction-time analyses included all remaining trials.

#### Pupillometry data exclusion

Three participants were excluded from all pupil-dilation analyses (one due to loss of signal across blocks, and two for whom eye-tracking recording was precluded by glasses). The remaining 29 participants entered the pupil analyses. For each participant, trials were excluded if more than 30% of samples within the analysis window (0 to 2 s post-stimulus) were lost to blinks or signal dropout.

#### Gaze data exclusion

One additional participant, beyond the three already excluded from pupil-dilation analyses, was excluded from gaze-based analyses due to insufficient valid fixation data (54.76% of trials contained at least two fixations within the stimulus region, below the 70% retention threshold), The remaining 28 participants were retained for gaze reinstatement and cumulative-fixation analyses. Within retained participants, trials with fewer than two fixations within the region of analysis were excluded from gaze similarity analyses. Final sample sizes varied by analysis according to the availability of valid trials for each comparison and were reported with each result.

### Data analysis

#### Behavioral measures

Accuracy in the one-back task was computed as the proportion of trials on which the observed response matched the correct response. *Same* trials (A-A) were trials with repeated items and required a *same* response. *Similar* trials were trials where the similar B item followed the A version (A-B trials) and required a *similar* response. *New* trials required no response. Response time was defined as the interval from stimulus onset to key press and was computed as the median response time across correct responses only.

Accuracy in the two-back task was computed separately for *same*, *similar* and *new* trials from each of the three conditions: (1) Items that had been *compared* to each other in the one-back phase, (2) items that had been encountered via *isolated* presentation, and (3) items that were *novel* in the two-back phase. *Similar* trials were those A-X-B trials requiring a “similar” response. *Same* trials were A-X-A trials requiring a “same” response. New trials were A-X-N trials requiring no key press. Response time was defined as the interval from stimulus onset to key press and was summarized as the median response time across correct trials only. A discrimination index was computed as the proportion of similar responses on similar trials minus the proportion of similar responses on new trials, following the standard defined in the Mnemonic Similarity Task as the lure discrimination index [22]. Same-item sensitivity was quantified using the signal detection measure *d^′^* [50], computed separately for each participant as *d^′^*= *z*(HR) *− z*(FAR), where *z*(*·*) denotes the inverse of the standard normal cumulative distribution function. The hit rate (HR) was the proportion of A-X-A trials receiving a “same” response, and the false alarm rate (FAR) was the proportion of A-X-N trials receiving a “same” response. To avoid undefined values arising from rates of 0 or 1, HR and FAR were bounded to the range [1*/*(2*N* ), 1 *−* 1*/*(2*N* )] before transformation, where *N* is the number of trials contributing to the respective rate (A-X-A trials for HR, A-X-N trials for FAR).

Recognition memory was measured separately for compared and isolated items in the final old-new recognition test. The hit rate (HR) was the proportion of old trials with a correct old response and response times greater than 150 ms. The false alarm rate (FAR) was the proportion of new foil trials with an old response and response times greater than 150 ms. Recognition sensitivity was computed as *d^′^* = *z*(HR) *− z*(FAR), where *z*(*·*) denotes the inverse of the standard normal cumulative distribution function; the same above described rate correction was applied before z transformation.

#### Gaze entropy

Spatial entropy was computed from normalized fixation density maps. Fixations were first restricted to the stimulus-centered region of interest (400 × 400 pixels spanning x: 760-1160, y: 340-740, corresponding to the 8.0° × 8.0° visual angle subtended by each stimulus). Each fixation map was divided into a 20 × 20 grid of spatial bins and normalized such that bin values summed to one. Entropy [51] was computed as *H* = *−* Σ*_i_ p_i_*log_2_ *p_i_*, where *p_i_* is the proportion of total fixation duration in bin i. Larger values indicate more distributed gaze patterns, whereas smaller values indicate more concentrated gaze patterns.

#### Episodic benefit

The episodic benefit was computed for each participant by comparing their performance on similar trials in the two-back phase between conditions. Specifically, the average discrimination index was contrasted between the items that had been encountered in the one-back phase (*compared* and *isolated* conditions) and items that were first encountered in the two-back phase (*novel* condition). Positive values indicate better discrimination for previously encountered pairs.

#### Pupil preprocessing

Pupil time series were preprocessed using the Pupil Common Drive Model (PCDM) toolbox [52]. Trial-level pupil data were first concatenated into a single continuous signal for each participant to avoid filter-edge artifacts. Each trial contributed a baseline segment from the 200 ms before stimulus onset and a stimulus-locked segment from stimulus onset to trial offset. Missing samples were set to zero before blink detection. Blinks were identified by thresholding the absolute sample-to-sample change in pupil diameter at one-tenth of the sampling rate. Detected blinks were removed and interpolated using the PCDM blink interpolation routine with margins of 5 samples before and 4 samples after each blink and interpolation windows of 50 and 75 ms. Remaining non-finite values were replaced by linear interpolation with nearest-neighbour extrapolation at signal boundaries. The concatenated signal was bandpass filtered between 0.03 and 10 Hz using a second order Butterworth filter and then segmented back into trial epochs. Participants with fewer than 30% valid samples across the concatenated signal were excluded from pupil analyses (see above).

#### Fixation density maps

Gaze analyses were performed on fixation density maps constructed within a stimulus-centred region of interest spanning x-coordinates 760-1160 pixels and y-coordinates 340-740 pixels on the 1920 × 1080 pixel display. Fixations outside this region were excluded. Remaining fixations were binned into a 20 × 20 grid, with each fixation contributing its dwell duration (ms) to its bin, so that longer fixations carried proportionally greater weight. Maps were smoothed with a Gaussian kernel with *σ* = 2 bins and normalized to unit sum, yielding duration-weighted fixation density maps [27].

#### Gaze similarity

Gaze similarity between two trials was defined as the Pearson correlation between their vectorized fixation density maps. To confirm that gaze similarity tracks item similarity, six types of comparison were computed separately for all items that appeared in both phases of the task (compared and isolated conditions): Same-item similarity measured gaze similarity for the same exemplar either across phases (A1-A2 similarity and B1-B2 similarity), or within phases (A1-A1 similarity and A2-A2 similarity). Similar-item similarity measured gaze similarity between perceptually similar but non-identical items either across phases (A2-B1 similarity and A1-B2 similarity), or within phases (A1-B1 similarity and A2-B2 similarity).

Additionally, because raw correlations between fixation density maps are inflated by viewing tendencies shared across trials (e.g., central bias, edge avoidance), for each participant a baseline-similarity was computed from combinations between all other (non-matching) B/A items within a given condition. This baseline was matched in participant, condition, and comparison type, such that any non-specific viewing tendencies are subtracted out symmetrically and cannot bias differences between conditions.

A gaze similarity index was then defined as the correlation of matching trials minus the average baseline correlation in that condition. Positive values indicate item-specific gaze similarity above the level expected from general spatial viewing tendencies.

#### Cumulative gaze similarity

To examine the time course of gaze similarity within two-back trials, correlations were recomputed cumulatively over successive fixations at A2. For each pair, the fixation density map was recomputed yielding updated fixation maps for up to four fixations (compare Supplementary Figure S4e). Because increasingly less trials were available for large numbers of fixations, a four-fixation limit was used; including a fifth fixation would have reduced the contributing sample below 20% of trials. At fixation steps 1-4, the proportion of trials contributing was 100%, 100%, 80%, and 48%, respectively. At each fixation step, the cumulative two-back map was then correlated with the full one-back map and baseline corrected using the same procedure described above. Cumulative similarity scores were averaged within participant at each fixation step and linear slopes across fixation steps were estimated for each participant.

### Statistical analysis

All statistical analyses were conducted on participant-level summary metrics unless otherwise specified. Repeated-measures models were used for all within-participant comparisons. Pairwise comparisons were conducted with paired-samples t-tests. Correction for multiple comparisons was realized by controlling the false discovery rate at *q* = 0.05 [53]. Tests of directional predictions specified a priori – the graded ordering of the episodic benefit across conditions (*compared > isolated > novel* ), the correct-versus-incorrect gaze and pupil contrasts, the cumulative-slope tests against zero, and the WM–EM correlations – were evaluated one-tailed; contrasts for which no directional prediction was made (the same-item control analyses and the compared-versus-isolated recognition comparison) were two-tailed. Repeated-measures ANOVAs were computed on participant-level cell means (MATLAB fitrm/ranova). Degrees of freedom are reported uncorrected; for effects with more than one numerator degree of freedom, *p*-values were Greenhouse–Geisser corrected and the correction factor *ε* is reported [54]. Effect sizes are reported as partial 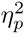 for ANOVA effects and as Cohen’s d for paired comparisons. Cohen’s d was computed as the mean within-participant difference divided by the standard deviation of that difference. In the analysis of behavioral responses and pupil dilation, Bayes factors were computed for the main repeated-measures condition effects using Bayesian ANOVA models with participant treated as a random factor [55].

#### Behavioral analyses

To test whether EM created in one-back affected WM performance in the subsequent two-back phase, we fit separate one-way repeated-measures ANOVAs for each behavioral variable (compare: Figure 2a-d). The model included encoding condition as a three-level within-participant factor (*compared*, *isolated* and *novel* ). The same ANOVA model was used to test for (1) mnemonic discrimination, (2) the d’ on same items, (3) the median reaction time on correct similar trials and (4) median reaction time on correct same trials. The model can be written as

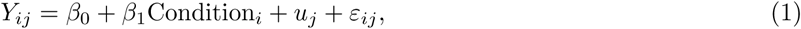

where *i* indexes condition and *j* indexes participant. Follow-up t-tests compared the *compared* and *isolated*, the *isolated* and *novel*, and the *compared* and *novel* conditions directly.

To test the association between memory systems and EM benefits across participants, (Figure 2e and Figure 2f), we computed Pearson correlations. Specifically, the episodic benefit for each participant was correlated with mean two-back same-item *d^′^*and with the recognition *d^′^*from the final old-new recognition test.

To assess task performance in the one-back phase, accuracy was compared across “same”, “similar” and “new” trials with paired-sample t-tests (Supplementary Figure S1). Additionally, median reaction time was compared between the correctly identified same and similar trials with a paired-samples t-test. Error patterns were summarized in a 3 *×* 3 confusion matrix as the proportion of realized responses (“same”, “similar”, “new”) that corresponded to each expected response type (“same”, “similar”, “new”).

#### Analyses of pupil dilation

To identify the time window in which pupil dilation differed by encoding condition, we performed separate cluster-based permutation tests across the post-stimulus time series for mnemonic discrimination trials (similar response required) and same-detection trials (same response required, compare: Supplementary Figure S3). At each time point, we fit a one-way repeated-measures ANOVA with encoding condition as a within-participant factor with the three levels: *compared*, *isolated*, and *novel*. At each time point, we also computed the pairwise paired-samples *t*-tests for *compared* versus *isolated*, *isolated* versus *novel*, and *compared* versus *novel*. Contiguous time points exceeding the statistical threshold corresponding to the 95th percentile of the cumulative distribution of the test-statistic (*F* = 3.162, corresponding to *p* = 0.05, for the omnibus effect), were grouped into clusters. This was done separately for the omnibus condition effect and for each pairwise contrast. Cluster statistics were then computed as the summed test statistic within each cluster and null distributions were generated by permuting condition labels within participant and retaining the maximum cluster statistic from each permutation. Finally, cluster-level *p*-values were computed as the proportion of permuted maximum cluster statistics that exceeded the observed cluster statistic.

Mean pupil dilation was then averaged within the significant time window from the omnibus test and analyzed with a post-hoc 3*×*2 repeated-measures ANOVA. This new model included encoding condition and discrimination accuracy as within-participant factors. Encoding condition had three levels: *compared*, *isolated*, and *novel*, while discrimination accuracy had two levels: *correct* and *incorrect*. The model can be written as

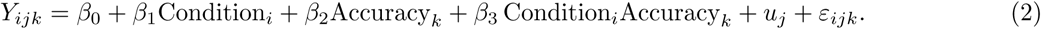

where *i* indexes condition and *j* indexes participant and *k* indexes accuracy (correct vs. incorrect). Follow-up t-tests compared encoding conditions and correct versus incorrect trials within each condition directly.

#### Gaze similarity analyses

To test whether fixation patterns carried item-specific information, baseline-corrected gaze similarity was averaged within participants for each item-comparison of interest. Same-item across-phase similarity was computed from A1-A2 and B1-B2 comparisons. Similar-item across-phase similarity was computed from A2-B1 and A1-B2 comparisons, and similar-item within-phase similarity was computed from A1-B1 and A2-B2 comparisons. Each measure was compared against a permutation-based null distribution generated by shuffling pair assignments within participants.

To test whether contrasting gaze patterns during one-back predicted later discrimination (pattern separation), the dependent variable A1-B1 gaze similarity was analyzed with a 2 × 2 repeated-measures ANOVA with encoding condition (isolated vs. compared presentation) and B2 mnemonic discrimination accuracy (correct vs. incorrect) as within-participant factors. To test if recapitulation of gaze patterns marked successful retrieval (pattern completion), B2-B1 gaze similarity was analyzed with the same model. To test whether anticipatory reinstatement during two-back predicted later discrimination (predictive recall), A2-B1 gaze similarity was analyzed with the same 2 × 2 repeated-measures ANOVA. Planned comparisons then tested correct versus incorrect trials within each encoding condition.

#### Cumulative gaze analyses

Cumulative reinstatement was tested with a 2 *×* 2 repeated-measures ANOVA on participant-level slopes. The model included similarity type and encoding condition as within-participant factors. Similarity type had two levels: A2–B1 and A2–A1. Encoding condition had two levels: *compared* and *isolated*. The model can be written as

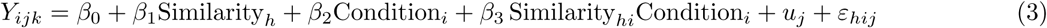

where *h* indexes similarity comparison, *i* indexes condition and *j* indexes participant. In a follow-up we tested and compared the slopes of the *compared* and *isolated* conditions separately. To this end, a linear slope was fitted across the cumulative gaze similarity measure. Trial-level slopes were then averaged for each participant, encoding condition, and similarity type and slopes were tested against zero with one-sample *t*-tests. Condition differences in slope were tested with paired-samples *t*-tests.

#### Spatial entropy analyses

To compare spatial entropy between conditions (Supplementary Information) a 2 × 2 repeated-measures ANOVA was used. The model included item position and encoding condition as within-participant factors. Item position had two levels (A and B), encoding condition had two levels (*compared* and *isolated* ). Follow-up t-tests compared conditions at each item position and compared item-positions within each condition

### Large language model use

The authors used a large language model (Claude Opus 4.8, Anthropic) to audit and review analysis code and for copyediting of the manuscript text. The model did not generate analyses, results, or scientific content. The authors take full responsibility for the work.

## Acknowledgements

We thank Jonathan Winawer, Clay E. Curtis, and Cate Hartley for helpful discussions, Jonathan Nicholas, Ricardo Alejandro, Leonard Niekerken, Chong Zhao, and Nikita Sossounov for helpful comments.

## Ethics approval

All procedures were approved by the New York University Institutional Review Board (protocol: FY2025-10338). The authors declare no competing interest.

## Data availability

Group statistical data that underly the analyses in this manuscript are available on zenodo (doi: 10.5281/zenodo.22017191). The individual data that support the findings of this study are available from the corresponding author upon request.

## Code availability

Analysis scripts that can produce the key figures and statistics in the manuscript are available on zenodo (doi: 10.5281/zenodo.22017336).

## Supplementary Information

**Supplementary Fig. S1.**
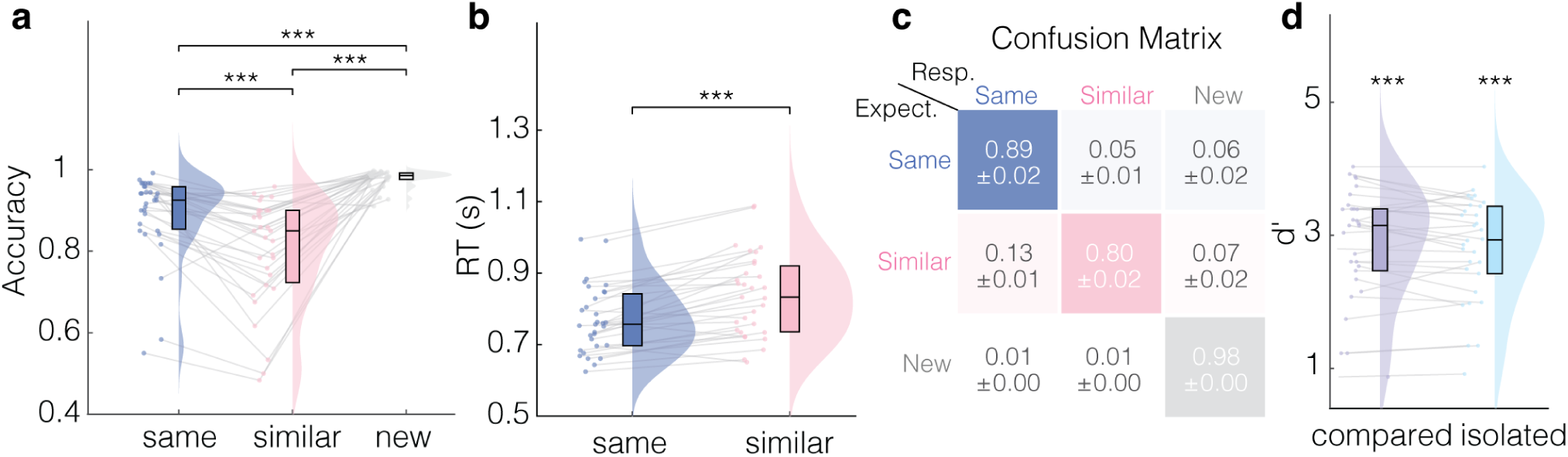
One-back task and recognition test performance. **a**, Accuracy in the one-back encoding task for same, similar, and new trials. Participants performed with high accuracy across all required response types (*same*: mean = 89%, SD = 10.4%; *similar* : mean = 80%, SD = 13.1%; *new* : mean = 98%, SD = 2.2%). Detection of identical repeats exceeded discrimination of similar items (*t*(30) = 5.52, *p < .*001, Cohen’s *d* = 0.99). Rejection of new items exceeded both *same* detection (*t*(30) = 4.44, *p < .*001, Cohen’s *d* = 0.80) and *similar* discrimination accuracy (*t*(30) = 7.49, *p < .*001, Cohen’s *d* = 1.35). **b**, Response time for correct same and similar trials during one-back encoding. Correct *same* responses were faster than correct *similar* responses (*t*(30) = *−*6.53, *p < .*001, Cohen’s *d* = 1.17). **c**, Confusion matrix for one-back encoding responses. Rows indicate the expected response category and columns the participant response; cells show mean response proportions *±* SEM. Correct responses dominated each row (*same* 0.89, *similar* 0.80, *new* 0.98); the most frequent error was misclassifying *similar* items as *same* (0.13). **d**, Recognition sensitivity (*d^′^*) for *compared* and *isolated* items in the final old/new recognition test. Both conditions were reliably above chance (*compared* : mean *d^′^* = 2.88, SD = 0.84, *t*(29) = 18.88, *p < .*001, *d* = 3.45; *isolated* : mean *d^′^*= 2.81, SD = 0.77, *t*(29) = 19.96, *p < .*001, *d* = 3.64) and did not differ from each other (*t*(29) = 1.53, *p* = .138, *d* = 0.28). Dots show individual participants, gray lines connect repeated measures, rainclouds show distributions, and box plots show medians and interquartile ranges. Asterisks indicate significance (*^∗∗∗^p < .*001).

**Supplementary Fig. S2.**
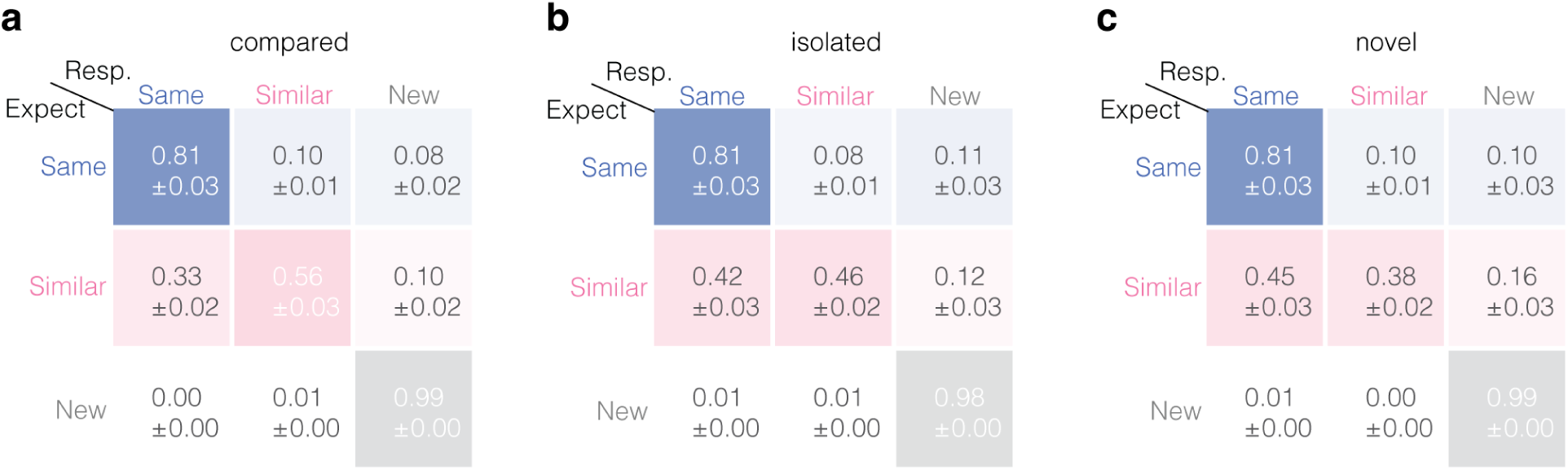
Two-back confusion matrices for similar-item trials. Confusion matrices for two-back retrieval trials that contained similar-item pairs, shown separately for **a**, *compared*, **b**, *isolated*, and **c**, *novel* conditions. Rows indicate the expected response category and columns the participant response; cells show mean response proportions *±* SEM. Correct *similar* responses were most frequent in the *compared* condition (0.56), lower in the *isolated* condition (0.46), and lowest in the *novel* condition (0.38), mirroring the graded discrimination index in Figure 2a. Across all three conditions the dominant error was misclassifying *similar* items as *same* (*compared* 0.33, *isolated* 0.42, *novel* 0.45); misclassification as *new* was comparatively rare (0.10–0.16).

**Supplementary Fig. S3.**
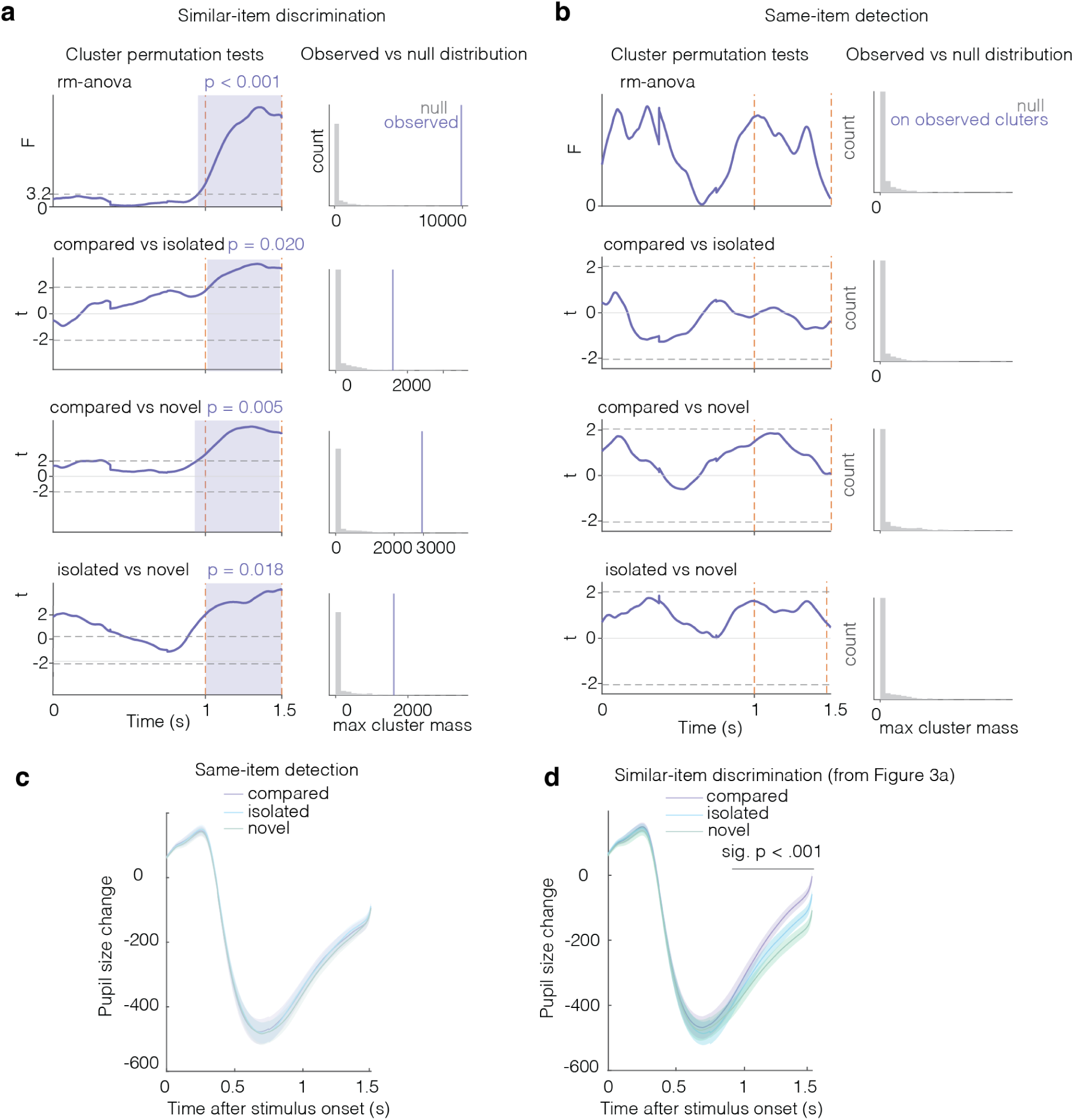
Pupil time-window selection and same-item control analysis. **a**, Cluster-based permutation tests for pupil dilation on *similar* -discrimination trials. The top row shows the omnibus effect of encoding condition (repeated-measures ANOVA) over time after stimulus onset; the lower three rows show the pairwise contrasts *compared* vs. *isolated*, *compared* vs. *novel*, and *isolated* vs. *novel*. Orange dashed lines mark the tested window and shaded regions mark significant clusters. A single late cluster (1.0–1.5 s) survived correction for the omnibus effect (*p < .*001) and for every pairwise contrast (*compared* vs. *isolated* : *p* = .020; *compared* vs. *novel* : *p* = .005; *isolated* vs. *novel* : *p* = .018); this window was used to summarize pupil change in Figure 3. Histograms on the right show the permutation null distributions of maximum cluster mass, with the observed cluster mass marked in purple. **b**, Corresponding cluster-based permutation tests for *same*-detection trials. No observed cluster exceeded the permutation null for the omnibus effect or for any pairwise contrast, indicating no reliable modulation of pupil dilation by encoding condition during same detection. **c**, Mean pupil time courses for *same*-detection trials across *compared*, *isolated*, and *novel* conditions. The overlapping traces are consistent with the null result in b and with the absence of a condition effect on averaged pupil change within the 1.0–1.5 s window (*F* (2, 56) = 1.00, *p* = .373, 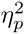 = .034, *ε* = .959, BF_01_ = 19.26). **d**, Mean pupil time courses for *similar* -discrimination trials, shown again from Figure 3a to allow direct comparison with same-item detection in c; the gray bar marks the 1.0–1.5 s cluster of significant modulation by condition (*p < .*001). Shaded regions in **c** and **d** show *±* SEM across *n* = 29 participants.

**Supplementary Fig. S4.**
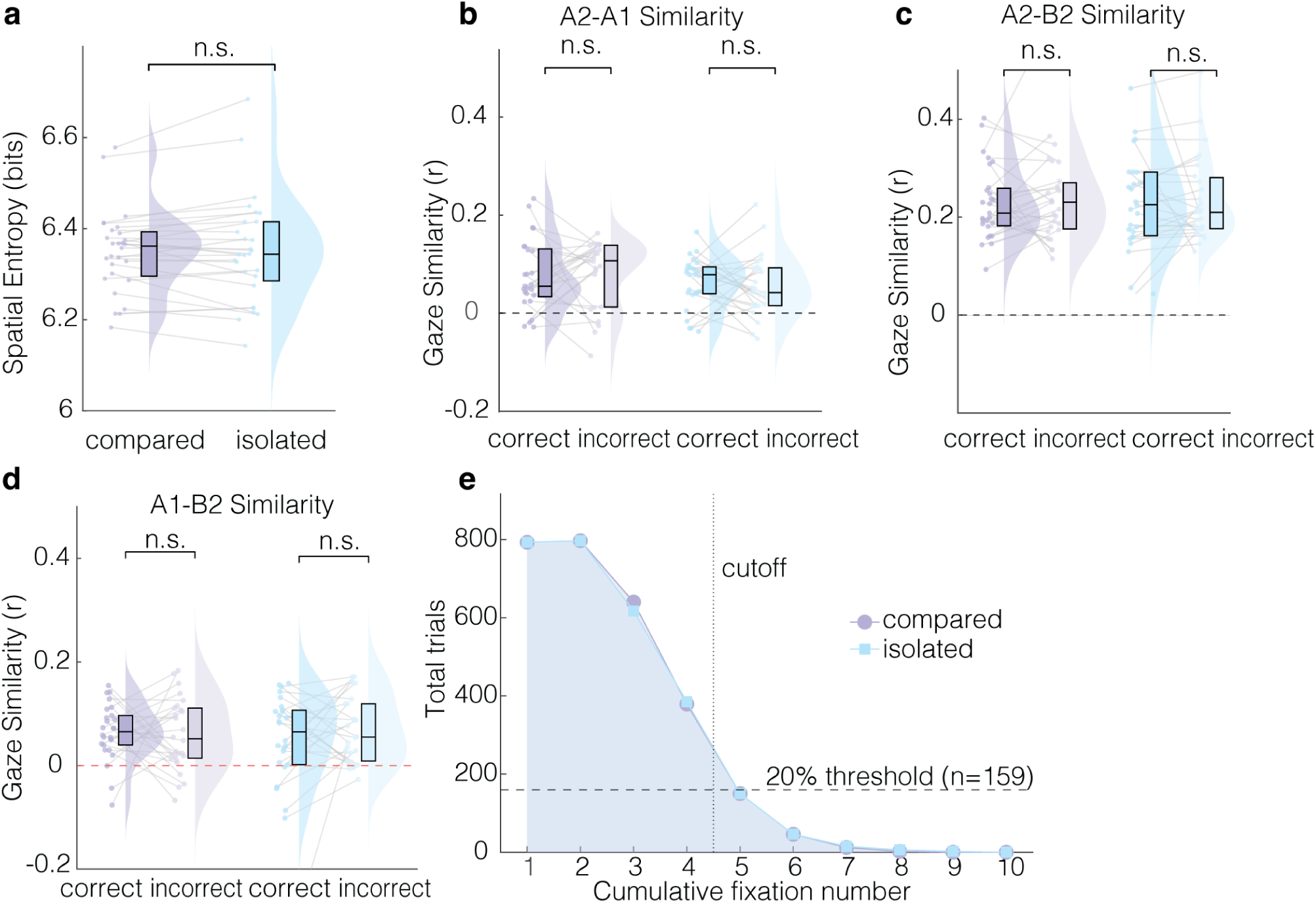
Gaze control analyses and trial counts for the cumulative-fixation analysis. **a**, Spatial entropy of fixation distributions for *compared* and *isolated* trials. A 2 *×* 2 (pair member [A/B] *×* encoding condition) repeated-measures ANOVA showed a main effect of pair member (*F* (1, 27) = 10.12, *p* = .004, 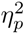 = .273; A items slightly higher than B) but no effect of encoding condition (*F* (1, 27) = 0.08, *p* = .779, 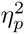 = .003, BF_01_ = 7.89) and no interaction (*F* (1, 27) = 0.30, *p* = .590, 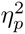 = .011), indicating that the condition differences in gaze similarity (Figure 4) were not explained by broader or more dispersed viewing. **b–d**, Control gaze-similarity analyses, each tested with the same 2 *×* 2 (encoding condition *×* subsequent B2 accuracy) ANOVA as the reinstatement measures in Figure 4d–f and each predicted to be null. **b**, A2–A1 gaze similarity indexes recapitulation of the A item’s own one-back viewing pattern. Discrimination accuracy did not modulate it (accuracy: *F* (1, 24) = 0.00, *p* = .973, 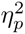 *< .*001; interaction: *F* (1, 24) = 0.39, *p* = .536, 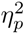 = .016; encoding condition: *F* (1, 24) = 3.34, *p* = .080, 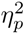 = .122; *compared* : *t*(24) = *−*0.28, *p* = .779, *d* = *−*0.06; *isolated* : *t*(24) = 0.40, *p* = .695, *d* = 0.08), confirming that predictive recall (A2–B1; Figure 4f) reflects anticipation of the counterpart rather than re-viewing of A. **c**, A2–B2 gaze similarity indexes how similarly the two pairmates perceived during two-back were viewed, and should not track memory; as expected it did not vary with accuracy (encoding condition: *F* (1, 26) = 0.08, *p* = .783, 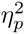 = .003; accuracy: *F* (1, 26) = 0.08, *p* = .785, 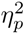 = .003; interaction: *F* (1, 26) = 0.38, *p* = .546, 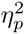 = .014; *compared* : *t*(26) = *−*0.66, *p* = .513, *d* = *−*0.13; *isolated* : *t*(26) = 0.23, *p* = .818, *d* = 0.04), showing that the reinstatement effects were not driven by generic perceptual similarity between pairmates at retrieval. **d**, A1–B2 gaze similarity indexes a non-diagnostic cross-phase pairing (A’s encoding with B’s retrieval) that does not correspond to reinstatement of B’s own encoding pattern; it too was unaffected by accuracy (encoding condition: *F* (1, 25) = 0.41, *p* = .528, 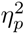 = .016; accuracy: *F* (1, 25) = 0.63, *p* = .434, 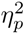 = .025; interaction: *F* (1, 25) = 1.35, *p* = .256, 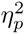 = .051; *compared* : *t*(25) = 0.24, *p* = .814, *d* = 0.05; *isolated* : *t*(25) = *−*1.23, *p* = .232, *d* = *−*0.24), confirming that pattern completion (B2–B1; Figure 4e) is specific to reinstatement of the counterpart’s encoding trace. **e**, Trial counts contributing to the cumulative-fixation analysis (Figure 5) as a function of cumulative fixation number; counts declined as more fixations were required, and the cutoff at four fixations marks the maximum retained, set by the 20% trial-count threshold (*n* = 159 trials). Purple, *compared* ; blue, *isolated*. In **b** through **d**, dots show individual participants, gray lines connect participant values, rainclouds show distributions, and box plots show medians and interquartile ranges; n.s., not significant.

## References

[1] Baddeley, A. Working memory. Science 255, 556–559 (1992). URL 10.1126/science.1736359.

[2] Luck, S. J. & Vogel, E. K. The capacity of visual working memory for features and conjunctions. Nature 390, 279–281 (1997). URL 10.1038/36846.

[3] Fuster, J. M. & Alexander, G. E. Neuron activity related to short-term memory. Science 173, 652–654 (1971). URL 10.1126/science.173.3997.652.

[4] Constantinidis, C. et al. Persistent spiking activity underlies working memory. The Journal of Neuroscience 38, 7020–7028 (2018). URL 10.1523/JNEUROSCI.2486-17.2018.

[5] Oberauer, K. & Lewandowsky, S. Further evidence against decay in working memory. Journal of Memory and Language 73, 15–30 (2014). URL 10.1016/j.jml.2014.02.003.

[6] Tulving, E. Elements of Episodic Memory Oxford Psychology Series (Oxford University Press, New York, 1983).

[7] Atkinson, R. & Shiffrin, R. Human Memory: A Proposed System and its Control Processes, 89–195 (Elsevier, 1968). URL 10.1016/S0079-7421(08)60422-3.

[8] Baddeley, A. D. & Hitch, G. Working Memory, 47–89 (Elsevier, 1974). URL 10.1016/S0079-7421(08)60452-1.

[9] Squire, L. R. & Zola-Morgan, S. The medial temporal lobe memory system. Science 253, 1380–1386 (1991). URL 10.1126/science.1896849.

[10] Oberauer, K. & Bartsch, L. M. When Does Episodic Memory Contribute to Performance in Tests of Working Memory? Journal of Cognition 6 (2023).

[11] Bartsch, L. M. & Oberauer, K. The contribution of episodic long-term memory to working memory for bindings. Cognition 231, 105330 (2023). URL 10.1016/j.cognition.2022.105330.

[12] Hoskin, A. N., Bornstein, A. M., Norman, K. A. & Cohen, J. D. Refresh my memory: Episodic memory reinstatements intrude on working memory maintenance. Cognitive, Affective, & Behavioral Neuroscience 19, 338354 (2019). URL 10.3758/s13415-018-00674-z.

[13] Yassa, M. A. & Stark, C. E. L. Pattern separation in the hippocampus. Trends in Neurosciences 34, 515–525 (2011).

[14] Bakker, A., Kirwan, C. B., Miller, M. & Stark, C. E. L. Pattern Separation in the Human Hippocampal CA3 and Dentate Gyrus. Science 319, 1640–1642 (2008).

[15] Favila, S. E., Chanales, A. J. H. & Kuhl, B. A. Experience-dependent hippocampal pattern differentiation prevents interference during subsequent learning. Nature Communications 7, 11066 (2016). URL 10.1038/ncomms11066.

[16] Michelmann, S. et al. Moment-by-moment tracking of naturalistic learning and its underlying hippocampo-cortical interactions. Nature Communications 12, 5394 (2021). URL 10.1038/s41467-021-25376-y.

[17] Hindy, N. C., Ng, F. Y. & Turk-Browne, N. B. Linking pattern completion in the hippocampus to predictive coding in visual cortex. Nature Neuroscience 19, 665–667 (2016).

[18] Horner, A. J. & Burgess, N. Pattern completion in multielement event engrams. Current biology : CB 24, 988–92 (2014). URL http://www.pubmedcentral.nih.gov/articlerender.fcgi?artid=4012134&tool=pmcentrez&rendertype=abstract.

[19] Horner, A. J., Bisby, J. A., Bush, D., Lin, W.-J. & Burgess, N. Evidence for holistic episodic recollection via hippocampal pattern completion. Nature Communications 6, 7462 (2015).

[20] Rolls, E. T. Pattern separation, completion, and categorisation in the hippocampus and neocortex. Neurobiology of Learning and Memory 129, 4–28 (2016).

[21] Ngo, C. T., Michelmann, S., Olson, I. R. & Newcombe, N. S. Pattern separation and pattern completion: Behaviorally separable processes? Memory & Cognition 49, 193–205 (2021).

[22] Stark, S. M., Kirwan, C. B. & Stark, C. E. Mnemonic similarity task: A tool for assessing hippocampal integrity. Trends in Cognitive Sciences 23, 938–951 (2019). URL 10.1016/j.tics.2019.08.003.

[23] Kirchner, W. K. Age differences in short-term retention of rapidly changing information. Journal of Experimental Psychology 55, 352–358 (1958). URL 10.1037/h0043688.

[24] Otero, S. C., Weekes, B. S. & Hutton, S. B. Pupil size changes during recognition memory. Psychophysiology 48, 1346–1353 (2011). URL 10.1111/j.1469-8986.2011.01217.x.

[25] Pajkossy, P., Szőllősi, Á. & Racsmány, M. Pupil size changes signal hippocampus-related memory functions. Scientific Reports 10, 16393 (2020). URL 10.1038/s41598-020-73374-9.

[26] Hannula, D. E. & Ranganath, C. The eyes have it: Hippocampal activity predicts expression of memory in eye movements. Neuron 63, 592–599 (2009). URL 10.1016/j.neuron.2009.08.025.

[27] Wynn, J. S., Shen, K. & Ryan, J. D. Eye movements actively reinstate spatiotemporal mnemonic content. Vision 3, 21 (2019). URL 10.3390/vision3020021.

[28] Wynn, J. S., Ryan, J. D. & Buchsbaum, B. R. Eye movements support behavioral pattern completion. Proceedings of the National Academy of Sciences 117, 6246–6254 (2020). URL 10.1073/pnas.1917586117.

[29] Mýzrak, E. & Oberauer, K. Working memory recruits long-term memory when it is beneficial: Evidence from the hebb effect. Journal of Experimental Psychology: General 151, 763–780 (2022). URL 10.1037/xge0000934.

[30] Langner, R. & Eickhoff, S. B. Sustaining attention to simple tasks: A meta-analytic review of the neural mechanisms of vigilant attention. Psychological Bulletin 139, 870–900 (2013). URL 10.1037/a0030694.

[31] Schacter, D. L., Dobbins, I. G. & Schnyer, D. M. Specificity of priming: a cognitive neuroscience perspective. Nature Reviews Neuroscience 5, 853–862 (2004). URL 10.1038/nrn1534.

[32] Kahneman, D. & Beatty, J. Pupil diameter and load on memory. Science 154, 1583–1585 (1966). URL 10.1126/science.154.3756.1583.

[33] van der Wel, P. & van Steenbergen, H. Pupil dilation as an index of effort in cognitive control tasks: A review. Psychonomic Bulletin & Review 25, 2005–2015 (2018). URL 10.3758/s13423-018-1432-y.

[34] Goldinger, S. D. & Papesh, M. H. Pupil dilation reflects the creation and retrieval of memories. Current Directions in Psychological Science 21, 90–95 (2012). URL 10.1177/0963721412436811.

[35] Kafkas, A. & Montaldi, D. Familiarity and recollection produce distinct eye movement, pupil and medial temporal lobe responses when memory strength is matched. Neuropsychologia 50, 3080–3093 (2012). URL 10.1016/j.neuropsychologia.2012.08.001.

[36] Papesh, M. H., Goldinger, S. D. & Hout, M. C. Memory strength and specificity revealed by pupillometry. International Journal of Psychophysiology 83, 56–64 (2012). URL 10.1016/j.ijpsycho.2011.10.002.

[37] Morris, C. D., Bransford, J. D. & Franks, J. J. Levels of processing versus transfer appropriate processing. Journal of Verbal Learning and Verbal Behavior 16, 519–533 (1977).

[38] Chanales, A. J., Oza, A., Favila, S. E. & Kuhl, B. A. Overlap among spatial memories triggers repulsion of hippocampal representations. Current Biology 27, 2307–2317.e5 (2017). URL 10.1016/j.cub.2017.06.057.

[39] Hulbert, J. C. & Norman, K. A. Neural differentiation tracks improved recall of competing memories following interleaved study and retrieval practice. Cerebral Cortex 25, 3994–4008 (2015). URL 10.1093/cercor/bhu284.

[40] Lewis-Peacock, J. A., Drysdale, A. T., Oberauer, K. & Postle, B. R. Neural evidence for a distinction between short-term memory and the focus of attention. Journal of Cognitive Neuroscience 24, 61–79 (2012). URL 10.1162/jocn_a_00140.

[41] Rose, N. S. et al. Reactivation of latent working memories with transcranial magnetic stimulation. Science 354, 1136–1139 (2016). URL 10.1126/science.aah7011.

[42] Wolff, M. J., Jochim, J., Akyürek, E. G. & Stokes, M. G. Dynamic hidden states underlying working-memory-guided behavior. Nature Neuroscience 20, 864–871 (2017). URL 10.1038/nn.4546.

[43] Goldman-Rakic, P. Cellular basis of working memory. Neuron 14, 477–485 (1995). URL http://linkinghub.elsevier.com/retrieve/pii/0896627395903046. ISBN: 0896-6273.

[44] Stokes, M. G. ‘activity-silent’ working memory in prefrontal cortex: a dynamic coding framework. Trends in Cognitive Sciences 19, 394–405 (2015). URL 10.1016/j.tics.2015.05.004.

[45] Beukers, A. O., Buschman, T. J., Cohen, J. D. & Norman, K. A. Is activity silent working memory simply episodic memory? Trends in Cognitive Sciences 25, 284–293 (2021). URL 10.1016/j.tics.2021.01.003.

[46] Brainard, D. H. The psychophysics toolbox. Spatial Vision 10, 433–436 (1997). URL 10.1163/156856897X00357.

[47] Pelli, D. G. The videotoolbox software for visual psychophysics: transforming numbers into movies. Spatial Vision 10, 437–442 (1997). URL 10.1163/156856897X00366.

[48] Kleiner, M., Brainard, D. & Pelli, D. What’s new in Psychtoolbox-3? Perception 36 (2007).

[49] Cornelissen, F. W., Peters, E. M. & Palmer, J. The eyelink toolbox: Eye tracking with matlab and the psychophysics toolbox. Behavior Research Methods, Instruments, & Computers 34, 613–617 (2002). URL 10.3758/BF03195489.

[50] Macmillan, N. A. & Creelman, C. D. Detection Theory (Psychology Press, 2004). URL 10.4324/9781410611147.

[51] Shannon, C. E. A mathematical theory of communication. Bell System Technical Journal 27, 379–423 (1948). URL 10.1002/j.1538-7305.1948.tb01338.x.

[52] Burlingham, C. S., Mirbagheri, S. & Heeger, D. J. A unified model of the task-evoked pupil response. Science Advances 8, eabi9979 (2022). URL 10.1126/sciadv.abi9979.

[53] Benjamini, Y. & Hochberg, Y. Controlling the false discovery rate: A practical and powerful approach to multiple testing. Journal of the Royal Statistical Society. Series B (Methodological) 57, 289–300 (1995). URL http://www.jstor.org/stable/2346101.

[54] Abdi, H. in The Greenhouse–Geisser correction (ed.Salkind, N.) Encyclopedia of Research Design 544–548 (Sage, Thousand Oaks, CA, 2010).

[55] Rouder, J. N., Morey, R. D., Speckman, P. L. & Province, J. M. Default Bayes factors for ANOVA designs. Journal of Mathematical Psychology 56, 356–374 (2012).

